# A Single-Cell Transcriptomic Atlas of Symmetry Breaking Across Eutherian Mammals

**DOI:** 10.64898/2026.02.27.708463

**Authors:** L. González-Brusi, N. Martínez de los Reyes, P. Marigorta, L. Simpson, A. Toledano-Díaz, J. Santiago-Moreno, R. Alberio, P. Bermejo-Álvarez, P. Ramos-Ibeas

## Abstract

The mammalian body plan is established during symmetry breaking and gastrulation. In ungulates and primates, these events occur within a flat embryonic disc and coincide with the developmental window most susceptible to pregnancy loss. Here, we generated a single-cell RNA sequencing atlas of *in vivo* sheep embryos that defines the timing and molecular programs underlying lineage segregation, as well as the inter-lineage signalling mediating anterior visceral hypoblast specification (AVH) and primitive streak formation.

By integrating sheep data with stage-matched datasets from cow, pig, rabbit, mouse, and marmoset, we constructed a cross-species atlas of anterior-posterior axis formation and identified conserved anterior and posterior hypoblast markers. Despite major differences in embryo morphology and developmental tempo, core signalling pathways NODAL, WNT, BMP, and FGF were broadly conserved. However, species-specific features emerged, including divergent BMP and FGF ligand usage and distinct BMP sources: only the mouse relied on trophectoderm-derived BMPs, whereas in other mammals BMPs originated primarily from the hypoblast. Functional experiments showed that NODAL is dispensable for early epiblast specification but essential for maintaining epiblast and AVH survival during symmetry breaking in sheep. These findings establish sheep as a powerful model for understanding human peri-implantation development and improving stem cell-based embryo models.

## Introduction

The establishment of the mammalian body plan is initiated during symmetry breaking and gastrulation, which specify the anterior-posterior axis and give rise to the embryonic germ layers from the pluripotent epiblast. Most current knowledge of these processes derives from studies in the mouse model, where gastrulation occurs in a three-dimensional egg cylinder ^1,2^. In contrast to this developmental architecture, symmetry breaking and gastrulation occur within a flat embryonic disc (ED) in most mammals, including primates and ungulates ^3^. A notable common feature of primates and ungulates during this developmental window is the high incidence of embryonic loss ^3–6^, underscoring the importance of elucidating the molecular mechanisms operating during this critical period.

While species-specific differences in implantation strategies and extraembryonic lineages development have been described ^7^, a conserved feature across species is the formation of an anterior signalling centre within the visceral hypoblast or endoderm, prior to primitive streak (PS) formation. This region secretes inhibitors of the BMP, WNT, and NODAL pathways, thereby restricting PS differentiation to the posterior epiblast ^8–10^. Single-cell transcriptomic analyses of *in vivo* embryos have enabled high-resolution mapping of symmetry breaking and gastrulation in several species, including mouse ^11,12^, rabbit ^13,14^, pig ^15^, non-human primates ^16,17^, and partially in humans ^18,19^. Nevertheless, functional validation of transcriptomic observations remains particularly challenging in primates, where symmetry breaking and gastrulation occur after implantation. Although human embryo models generated from stem cells have enabled important advances in the field, they remain largely informed by mouse embryo-based frameworks, do not fully recapitulate the inter-lineage interactions present *in vivo*, and still offer limited utility for functional studies ^12,20^.

Ungulates provide an accessible model for studying symmetry breaking and gastrulation *in vivo*, as these processes occur while the embryo remains free-floating within the uterine lumen, during a phase known as conceptus elongation. In sheep, following hatching from the *zona pellucida*, the conceptus undergoes a prolonged preimplantation period characterized by exponential expansion of the extraembryonic lineages (trophectoderm and hypoblast) alongside the initiation of symmetry breaking and gastrulation within the ED between embryonic day (E) 11 and 14 ^21,22^.

To advance our understanding of symmetry breaking and gastrulation in mammals developing a flat ED, we present a single-cell transcriptomic atlas of *in vivo*-derived sheep embryos collected between E11 and E13.5. This dataset reveals the timing, lineage markers, and signalling pathways driving the emergence of key embryonic and extra-embryonic lineages. We further dissect the molecular mechanisms and inter-lineage communication underlying symmetry breaking across six mammalian species. Finally, by integrating transcriptomic and loss-of-function approaches, we demonstrate the essential role of NODAL signalling in epiblast and anterior visceral hypoblast (AVH) survival in sheep.

## Results

### A single-cell map of peri-gastrulation and conceptus elongation in sheep

To investigate the emergence and progression of embryonic and extraembryonic lineages during gastrulation and conceptus elongation, we performed single-cell RNA sequencing (scRNA-seq) on a total of 15,767 cells from eight pooled samples, comprising 80 *in vivo*-derived embryos collected between embryonic day (E) 11, prior to symmetry breaking and onset of gastrulation, and 13.5, after gastrulation but before neural folding and somitogenesis. The dataset included E11 and E11.5 whole embryos (spherical and ovoid), as well as isolated embryonic discs (EDs) with surrounding extraembryonic membranes (EEMs) from E12.5 and E13.5 (tubular and filamentous) conceptuses (**Fig. 1a, Supplementary Data 1**, **Supplementary Fig. 1**). After quality control, transcriptomes of 13,847 cells were retained, with a median of 4,704 genes detected per cell (**Supplementary Fig. 2a, b, c**).

**Figure 1.**
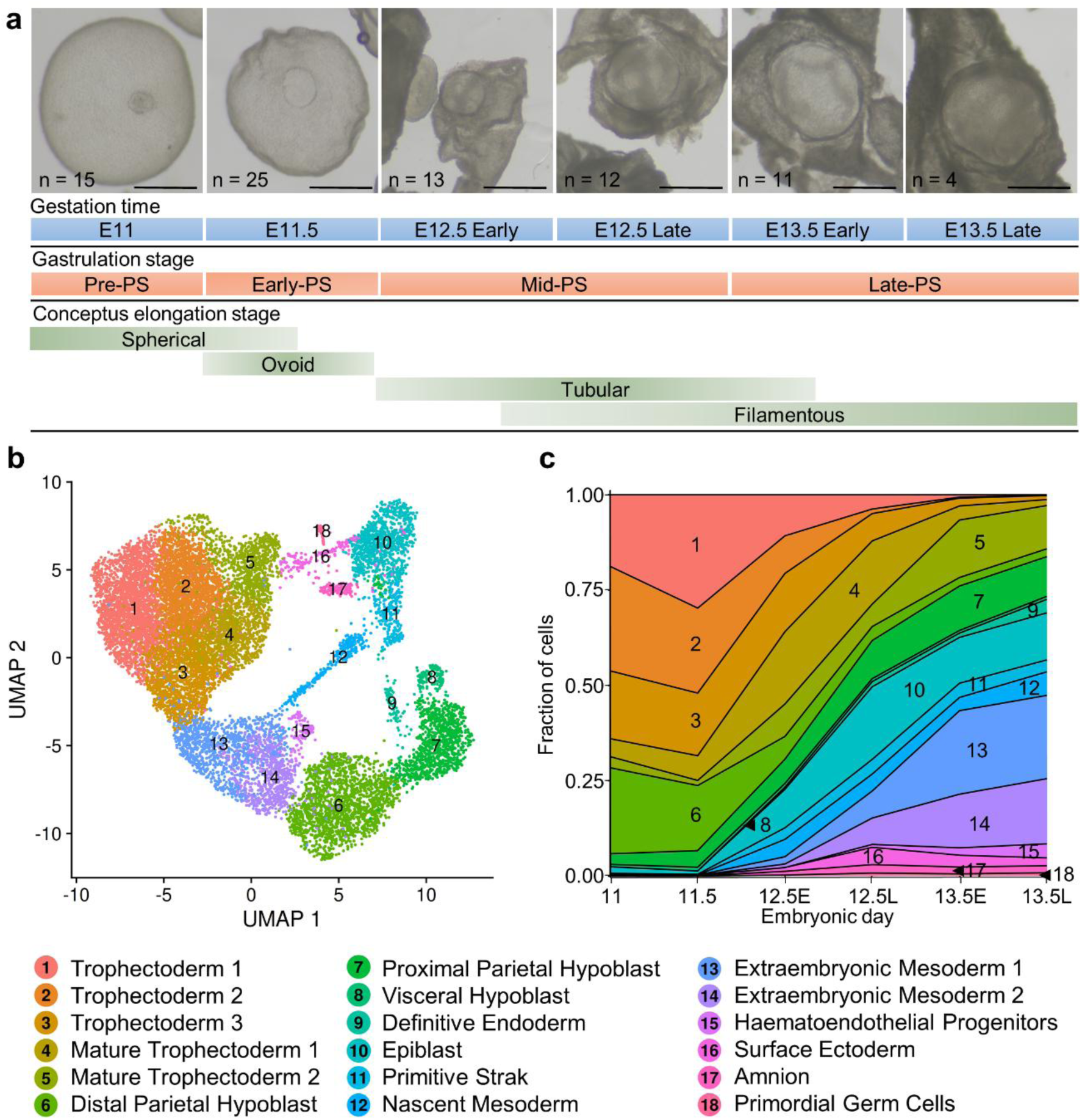
Sheep single-cell transcriptional atlas of conceptus elongation, symmetry breaking and gastrulation. a) Representative images of sheep embryos collected from embryonic day (E) 11 to 13.5, including whole embryos and isolated embryonic discs (EDs) with surrounding extraembryonic membranes (EEMs) (for sample collection details, See Supplementary Data 1). Scale bars: 500 µm. PS: primitive streak. b) Uniform manifold approximation and projection (UMAP) plot of 15,767 sheep cells from eight pooled embryo samples. Cells are coloured according to their cell-type annotation and numbered as indicated in the legend below. c) Stacked area plot displaying the fraction of each cell type across time points. The proportion of extraembryonic membranes (EEMs: trophectoderm and hypoblast) decreases in isolated embryonic discs (EDs), while cell-type diversity increases after E12.5, with mesodermal lineages becoming predominant in the ED.

Unsupervised clustering, integrated with developmental stage information and canonical lineage markers, was used to define 18 distinct cell clusters (**Fig. 1b, c**, **Supplementary Fig. 2d, e**, **Supplementary Data 2**), and enabled the identification of novel marker genes for specific cell populations in sheep (**Supplementary Data 3**). Spherical and ovoid embryo samples were mainly represented by extra-embryonic lineages: trophectoderm (TE) and hypoblast, whereas isolated EDs from E12.5 onward were enriched in embryonic lineages.

### Mapping the development of extraembryonic membranes in sheep

Five TE populations were identified (**Fig. 1b**). Immature TE1, TE2, and TE3 clusters showed high expression of *CDX2* and *LGALS3* (**Supplementary Fig. 3a, b**) and were more abundant in E11 and 11.5 spherical and ovoid embryos (**Figure 1c, Supplementary Fig. 2d**). In contrast, mature TE clusters, primarily composed of cells from E12.5 onward in tubular and filamentous conceptuses, exhibited high expression of *IFNT5, IFNT6,* and *IFNT11* (encoding interferon-tau, the main pregnancy recognition signal in ungulates ^23^), as well as *FURIN* and other previously reported mature TE markers such as *PTGS2* and *PAG11* ^24^. Interestingly, genes such as *FADS1, HAND1,* and *PINLYP*, previously associated with immature TE, were also upregulated in the mature TE population (**Supplementary Fig. 2e and 3a, b**). TE maturation in ungulates involves the emergence of binucleate cells (BNCs), which fuse with endometrial epithelial cells to form trinucleate cells after implantation ^25^. Although few BNCs were detected via immunostaining for GATA3 and F-actin membrane labelling in E13.5 embryos (**Supplementary Fig. 3c**), BNC-specific markers, including *CSH1, CSH2, PAG10, PAG16, PAG17, PRP1,* and *PRP2,* were not detected in our dataset, likely due to the scarcity of this population at this stage. Gene ontology (GO) analysis of differentially expressed genes (DEGs) revealed enrichment for biological processes such as active transmembrane transporter activity, estrogen response element binding, and phosphoric ester hydrolase activity in immature TE cells during early conceptus elongation. Mature TE cells, on the other hand, exhibited increased expression of genes involved in SMAD binding and lipid transmembrane transporter activity, among other functions (**Supplementary Data 4**).

Among the three hypoblast populations identified, distal parietal hypoblast (dPH) consisted of E11 and 11.5 cells from spherical and ovoid embryos, showing high expression of *PDGFRA, BMP7,* and *CRIPTO*. In contrast, proximal parietal hypoblast (pPH) cells mainly originated from isolated EDs with surrounding EEMs from E12.5 onward, and expressed *PRDM1*, *BMP2,* and *BMP6*, as well as genes involved in the biding of various molecules, including type II TGF-beta receptor, glycosaminoglycans, vitamins, or carboxylic acid, among others. The visceral hypoblast (VH) population showed specific expression of *NODAL, EOMES, OTX2, LHX1,* and *HHEX.* Within this cluster, a subset of cells also expressed *CER1, LEFTY1,* and *DKK1* (**Fig. 1b and c, Supplementary Fig. 2d, 2e, 3a, 3d, Supplementary Data 5**).

**Figure 2.**
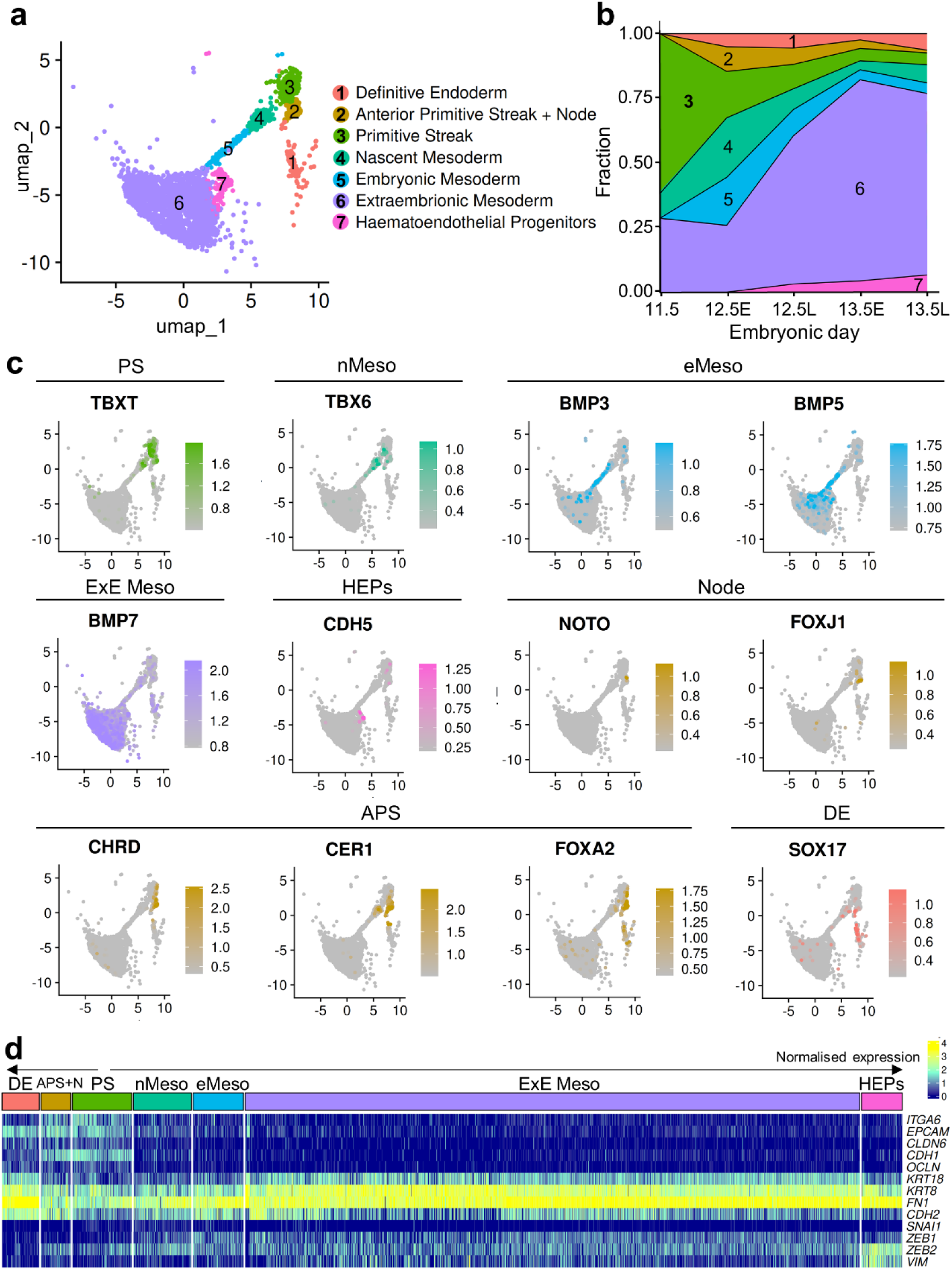
Rapid mesoderm progenitors differentiation and proliferation during early gastrulation. a) UMAP plot showing subclustering of primitive streak (PS), mesoderm, and definitive endoderm (DE) populations identified in Figure 1b (clusters 9, 10, 12, 13, 14, and 15). b) Stacked area plot displaying the temporal distribution of lineages in a). c) Feature plots coloured by normalised gene expression of selected marker genes. Feature plot scales were determined by the 5^th^ and 95^th^ percentiles. d) Heatmap showing the normalised expression of epithelial and mesenchymal marker genes within individual cells in subclusters from a). APS: anterior primitive streak; DE: definitive endoderm; eMeso: embryonic mesoderm; ExE Meso: extraembryonic mesoderm; HEPs: haematoendothelial progenitors; nMeso: nascent mesoderm; PS: primitive streak.

**Figure 3.**
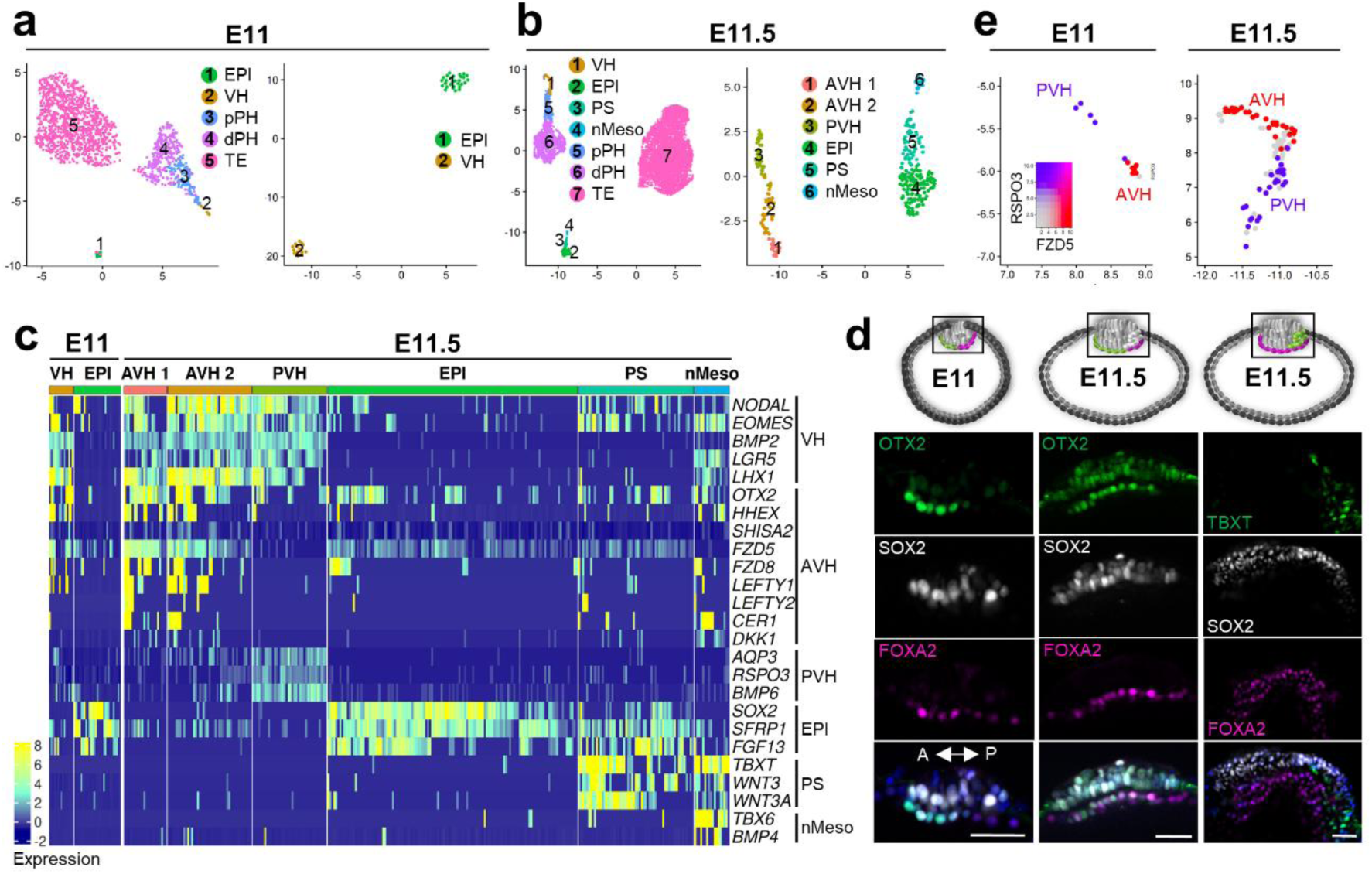
Onset of symmetry breaking in sheep embryos. a) Uniform manifold approximation and projection (UMAP) plots of all lineages present in E11 embryos, and subclustering of populations present within the ED: EPI and VH. b) UMAP of all lineages present in E11.5 embryos and subclustering of EPI, PS, nMeso, and VH populations. c) Heatmap showing the scaled expression of canonical and novel markers within individual cells in the subpopulations emerging from subclusterings in a) and b). d) Schemes depicting the lineages shown in the images. Representative sagittal z-stack images of E11 and E11.5 EDs stained for OTX2 (epiblast and visceral hypoblast), SOX2 (epiblast), FOXA2 (hypoblast) and TBXT (mesoderm). Scale bars: 50 µm. e) UMAPs of E11 and E11.5 VH cells showing unique *FZD5* and *RSPO3* expression patterns in anterior and posterior VH cells. AVH: anterior visceral hypoblast; ED: embryonic disc; EPI: epiblast; dP HYP: distal parietal hypoblast; nMeso: nascent mesoderm; pP HYP: proximal parietal hypoblast; PS: primitive streak; PVH: posterior visceral hypoblast; TE: trophectoderm; VH: visceral hypoblast.

### Rapid mesoderm differentiation and proliferation during early gastrulation in sheep

Primitive streak (PS), nascent mesoderm (nMeso), and extraembryonic mesoderm (ExEMeso) cells were detected from E11.5. From E12.5 onward, mesodermal populations expanded rapidly, whereas definitive endoderm (DE) cells exhibited slower proliferative dynamics. Surface ectoderm, amnion and primordial germ cells (PGCs) also emerged from E12.5 onwards (**Fig. 1b, c**).

To investigate early gastrulation events, we performed subclustering of the PS, mesoderm, haematoendothelial progenitors (HEPs) and DE clusters identified in the initial unsupervised clustering (**Fig. 1b**: clusters 9, 10, 11, 12, 13, 14, and 15). The PS, characterized by high *TBXT* expression, was detected from E11.5. A mixed population expressing anterior primitive streak (APS) markers (*CHRD*, *CER1*, and *FOXA2*) along with some cells expressing node markers (*NOTO* and *FOXJ1*) emerged in early E12.5 embryos and diminished as the DE became established. A *TBX6-*positive nascent mesoderm (nMeso) population was present at E11.5 and declined over time. *BMP7*-positive ExE Meso cells were also detected as early as E11.5, coincident with the appearance of the PS, and exhibited exponential expansion thereafter. *Bmp5* expression has been associated with presumptive anterior mesoderm in mice, whereas *Bmp7* exhibits a broader expression pattern ^26^. Notably, a distinct *BMP3-* and *BMP5*-positive embryonic mesoderm population emerged later, at E12.5. This population expressed genes indicative of anterior or embryonic mesoderm identity ^27,28^, precursor to lateral plate, intermediate, paraxial and axial mesoderm. However, these advanced mesoderm lineages were not distinctly resolved within the developmental stages analysed in our dataset, suggesting their emergence occurs at later stages, consistent with observations in human Carnegie Stage 8 embryos ^19^. The ExE Meso cluster was enriched in genes upregulated in the posterior mesoderm in mice (**Supplementary Figure 4**) ^27,28^. HEPs, marked by exclusive expression of *CDH5,* were first detected from late E12.5 onwards (**Fig. 2a-c**).

Consistent with previous findings in pig gastrulation ^15^, epithelial markers *ITGA6* and *CDH1* were downregulated in mesodermal cells emerging from the PS. Similarly, cell-cell adhesion-associated gene *EPCAM*, along with tight junction markers *OCLN* and *CLDN6,* were downregulated in mesodermal populations but maintained in APS, node, and DE. The intermediate filament genes *KRT8* and *KRT18*, as well as other genes involved in cell-cell adhesion, including *FN1* and *CDH2*, were upregulated in both emergent mesoderm populations and DE. Conversely, epithelial-to-mesenchymal transition (EMT) regulator *SNAI1* was upregulated in nMeso cells. *ZEB1* and *ZEB2*, transcriptional repressors of *CDH1*, were highly expressed in mesoderm populations, and the mesenchymal marker *VIM* was upregulated in HEPs (**Fig. 2d**). Together, these data suggest that APS, node and DE populations largely retain an epithelial identity in contrast to mesodermal derivatives of the PS, which undergo EMT, consistent with observations in the pig embryo ^15^.

DEGs analysis revealed upregulation of some VH-associated markers in the DE, such as *BMP2, LGR5,* and *AQP3*. In contrast, *NODAL, EOMES, OTX2* were enriched in PS and APS + Node subclusters (**Supplementary Data 6, Fig. 2c**). GO terms analysis revealed enrichment for chemorepellent activity in DE and APS + Node populations, as well as for semaphorin receptor binding in APS + Node, similar to what was detected in the AVH. These findings could be linked to the spatial and temporal proximity of these cell populations.

### Exploring the onset of symmetry breaking in sheep

The onset of anterior-posterior (A-P) polarity is initiated by the specification of a signalling centre in the anterior hypoblast, which prevents PS formation in the adjacent epiblast. The emergence of this anterior organizer, termed the anterior visceral endoderm (AVE) in the mouse egg cylinder or anterior visceral hypoblast (AVH) in species developing an ED, appears conserved across mammalians ^29,30^. In mice, a subset of hypoblast cells (known as primitive endoderm, PrE) at the distal tip of the embryo, in response to the diminished exposure to signals derived from the extraembryonic ectoderm (ExE; derived from the polar TE in direct contact with the epiblast) that include BMP4, gives rise to the distal visceral endoderm (DVE), which initiates expression of the NODAL antagonist *Lefty1*^30,31^. Whether the AVE arises directly from the DVE or instead from *Lefty1*^−^ cells within the visceral endoderm remains unresolved^12,32^, although in both scenarios, these cells migrate from a distal to an anterior position to establish the A-P axis. Nevertheless, the developmental origin and molecular pathways underlying AVH emergence in species where symmetry breaking occurs in a two-dimensional ED remain uncharacterized.

To investigate the molecular determinants of symmetry breaking in the sheep embryo, we focused on the ED by subclustering epiblast and *NODAL-, EOMES-,* or *OTX2*-positive visceral hypoblast (VH)-derived populations at the earlier stages in our dataset: E11 and 11.5. At E11, unsupervised clustering did not reveal additional cell populations (**Fig. 3a**), and no TBXT-positive cells were detected (**Fig. 3c**), indicating that PS formation had not yet started. Subclustering of EPI, PS, nascent mesoderm (nMeso), and VH cells from E11.5 identified six subpopulations: *SOX2*-positive EPI; PS cells downregulating *SOX2* and upregulating *TBXT*; *TBX6*-positive nascent mesoderm (nMeso); two AVH populations; and a VH population not expressing AVH markers, which we termed posterior VH (PVH) (**Fig. 3b, c**).

Most VH cells at both E11 and 11.5 expressed classical markers such as *NODAL* and *EOMES*, as well as *LGR5* and *LHX1,* previously reported as AVH markers in NHP and human embryos ^12,16^. *OTX2* was also broadly expressed in the VH, but enriched in AVH clusters compared to PVH at E11.5. Immunofluorescence confirmed this asymmetric distribution, along with weaker expression in EPI (**Fig. 3d**), a pattern conserved in mice ^33^, rabbits, pigs ^34^, and primates ^16^.

No clear molecular signature of the DVE was detected, characterized by low *LEFTY1* and high *IHH*, *NRP1*, and *DKK1*. DVE precursor markers such as *WNT3*, *FGF8,* and *FGF10* were not identified, in agreement with findings in human and non-human primates (NHP) ^12^. Similarly, no evidence of DVE formation has been reported in rabbits, where *DKK1, LEFTY1/2,* and *CER1* expression is initially diffuse across the VH and later becomes localised anteriorly ^34^. This suggests divergent mechanisms underlying initial symmetry breaking in species with a flat ED compared to the mouse-specific three-dimensional egg cylinder.

At E11, *NODAL, HHEX,* and *LEFTY1* were expressed in a VH subpopulation negative for *CER1* and *DKK1*, as previously reported in pig embryos prior to symmetry breaking ^34^, and consistent with an early AVH or pre-AVE identity as reported in mice ^12^. By E11.5, *CER1, LEFTY2* and *DKK1* expression emerged in AVH1 and AVH2 clusters, confirming a more mature AVH identity.

Additional AVH-specific markers, including *SHISA2, FZD5,* and *FZD8* ^12,16,35^ were absent in the PVH (**Fig. 3c**). DEG analysis further identified specific markers for the AVH1, AVH2, and PVH populations (**Supplementary Data 7**). Among these, *AQP8, SEMA6A,* and *NRG1*, early DVE/AVE markers in mice, were upregulated in AVH2. GO analysis of AVH2-specific genes revealed enrichment for chemorepellent activity, similar to what has been reported for the early mouse AVE ^12^. Together, these findings suggest that AVH2 may represent an earlier AVH population than AVH1, or possibly an intermediate state between AVH1 and PVH, consistent with the observation that the second UMAP dimension reflects A-P positioning of EPI, PS, and nMeso cells (**Fig. 3b**).

*AQP3* and *RSPO3*, previously detected in the mouse visceral endoderm ^12^, as well as *BMP6*, were specifically expressed in the PVH (**Figure 3c**). When mapping the WNT receptor *FDZ5* ^35^ and the WNT activator *RSPO3* ^36^ across all VH cells, FZD5^+^/RSPO3^−^ anterior and FZD5^−^/RSPO3^+^ posterior VH populations were evident at both E11 and E11.5 (**Fig. 3e**), indicating the emergence of a WNT signalling gradient within the VH as early as E11.

### Cross-species transcriptome atlas of symmetry breaking onset

To investigate the molecular mechanisms underlying symmetry breaking across mammals, we analyzed scRNA-seq datasets from cow ^37^, pig ^15,38^, rabbit ^14^, mouse ^39–41^, and marmoset ^16^. Equivalent embryonic stages and lineages were extracted to match sheep pre-PS (E11) and early-PS (E11.5) cell populations forming the ED, together with surrounding EEMs, including TE and proximal parietal hypoblast (PH). At the pre-PS stage, anterior and posterior VH clusters were already evident in cow and rabbit embryos, and the DVE/AVE could be distinguished from the remaining embryonic visceral endoderm (emVE) in mouse. In contrast, VH cells in sheep, pig and marmoset embryos did not segregate into anterior and posterior domains with asymmetric expression of classical markers (*LEFTY1*, *CER1*, *HHEX*, *DKK1*, *SHISA2*) (**Fig. 4b-g**). By the early-PS stage, AVH/AVE and PVH/emVE populations were detected in all species (**Fig. 4h-m**).

**Figure 4.**
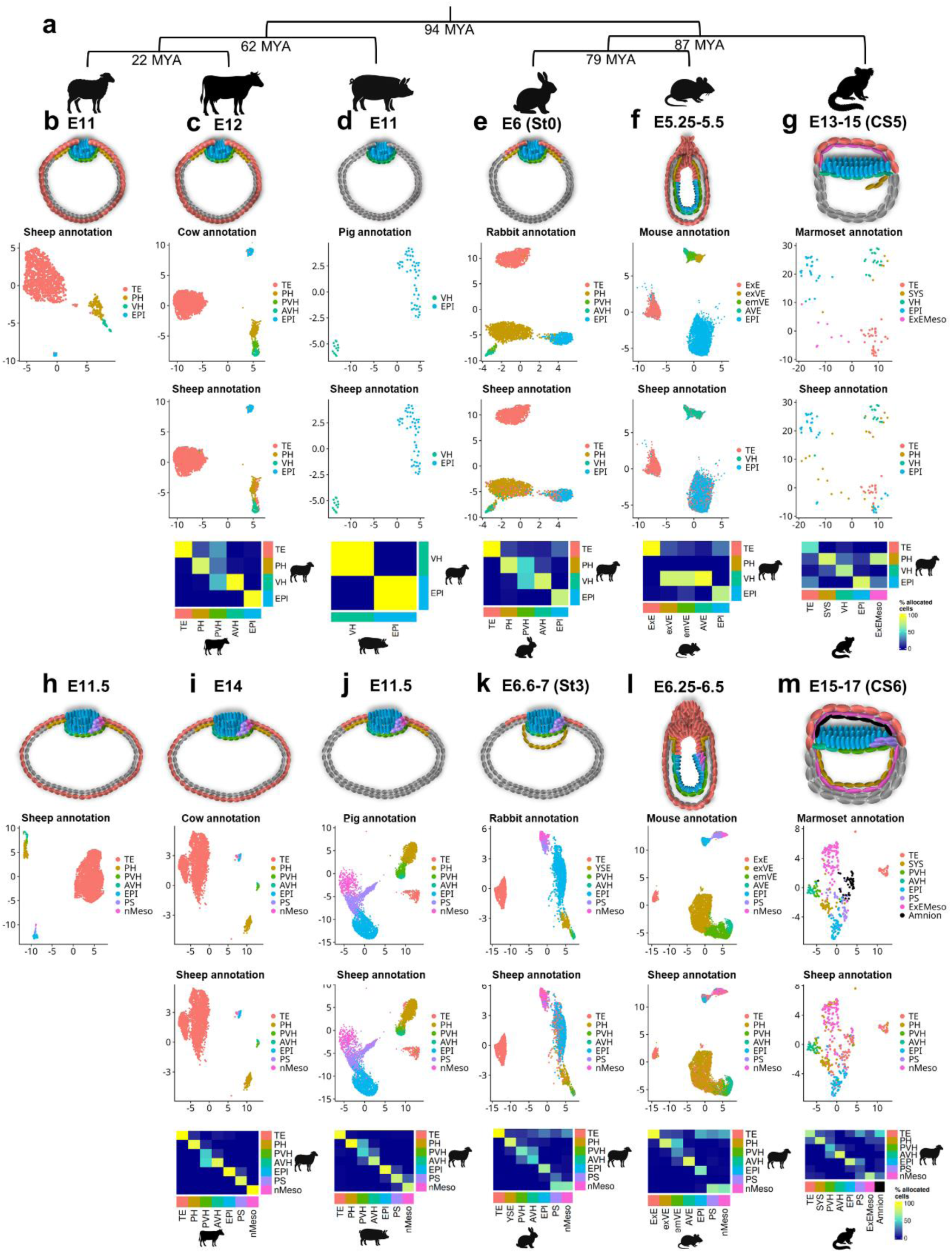
Alignment of sheep, cow, pig, rabbit, mouse, and marmoset datasets. a) Phylogenetic tree of sheep, cow, pig, marmoset, and mouse. Distances are not to scale; divergence times (in million years ago; MYA) are based on the Timetree tool ^43^. b-g) Schematic representation of the selected cell populations from b) E11 sheep (this dataset), c) E12 cow (*Bos Taurus*)^37^, d) E11 pig (*Sus scrofa)* embryos ^38^, e) E6 (Stage 0) rabbit (*Oryctolagus cuniculus*) embryos ^14^, f) E5.25-5.5 mouse (*Mus musculus*) embryos ^39–41^, or g) E13-15 (CS5) marmoset (*Callithrix jacchus*)^16^ pre-primitive streak (pre-PS) embryos, and Uniform Manifold Approximation and Projection (UMAP) plots showing selected cell populations, shown either with the original (cow/pig/rabbit/mouse/marmoset) annotation or with the sheep annotation after reciprocal PCA-based projection onto the respective dataset. The accompanying heatmap indicates the percentage of c) cow, d) pig, e) rabbit, f) mouse, and g) marmoset cells in each cluster allocated to the corresponding sheep cell populations after label transfer. h-m) Schematic representation of the selected cell populations from h) E11.5 sheep (this dataset), i) E14 cow ^37^, j) E11.5 pig embryos ^15^, k) E6.6-7 (Stage 3) rabbit embryos ^14^, l) E6.25-6.5 mouse embryos ^39^, or m) E15-17 (CS6) marmoset ^16^ early-primitive streak (early-PS) embryos, and UMAP plots showing selected cell populations from, shown either with the original (cow/pig/rabbit/mouse/marmoset) annotation or with the corresponding sheep annotation after reciprocal PCA-based projection onto the respective dataset. The heatmap displays the percentage of i) cow, j) pig, k) rabbit, l) mouse, and m) marmoset cells in each cluster allocated to the corresponding sheep cell populations after label transfer. AVH: anterior visceral hypoblast; EPI: epiblast; emVE: embryonic visceral endoderm; ExEct: extraembryonic ectoderm; ExEMeso: extraembryonic mesoderm; exVE: extraembryonic visceral endoderm; nMeso: nascent mesoderm; PH: parietal hypoblast; PS: primitive streak; PVH: posterior visceral hypoblast; SYS: secondary yolk sac; TE: trophectoderm; YSE: yolk sac endoderm.

Detection of anchor genes between sheep and cow, pig, rabbit, mouse, and marmoset datasets, followed by label transfer, revealed strong correlations between equivalent cell-type annotations. In marmoset, early differentiating extraembryonic mesoderm (ExEMeso) showed a high correlation with the sheep PH, consistent with the hypothesis that the earliest ExEMeso originates from the hypoblast in primates ^42^. The secondary yolk sac (SYS), a primate-specific structure emerging from the VH, and the rabbit yolk sac endoderm (YSE), also matched the sheep PH (**Fig. 4b-m**).

To identify highly conserved anterior and posterior markers across species, we compared the DEGs identified using *FindMarkers* on AVH/AVE and PVH/emVE clusters (**Fig. 5**). Canonical markers such as *CER1*, *DKK1*, *LHX1*, *HESX1,* and *HHEX* were consistently upregulated in the AVH, with *HHEX* being AVH-specific in all species. *OTX2, FZD5*, and *GSC* also displayed anterior specificity in most species, as in sheep. Additional anterior markers common to ungulates and rodents included *TCF7L2, SEMA6A, CRIM1, LRIG3,* and *ROR1.* Conserved posterior markers included *MSX1, FGB, FBLN2, SLC39A5,* and *RSPO3*, a potent WNT signalling enhancer ^36^ and a previously reported hypoblast marker in primates ^44^ that may contribute to WNT activation in the posterior EPI.

**Figure 5.**
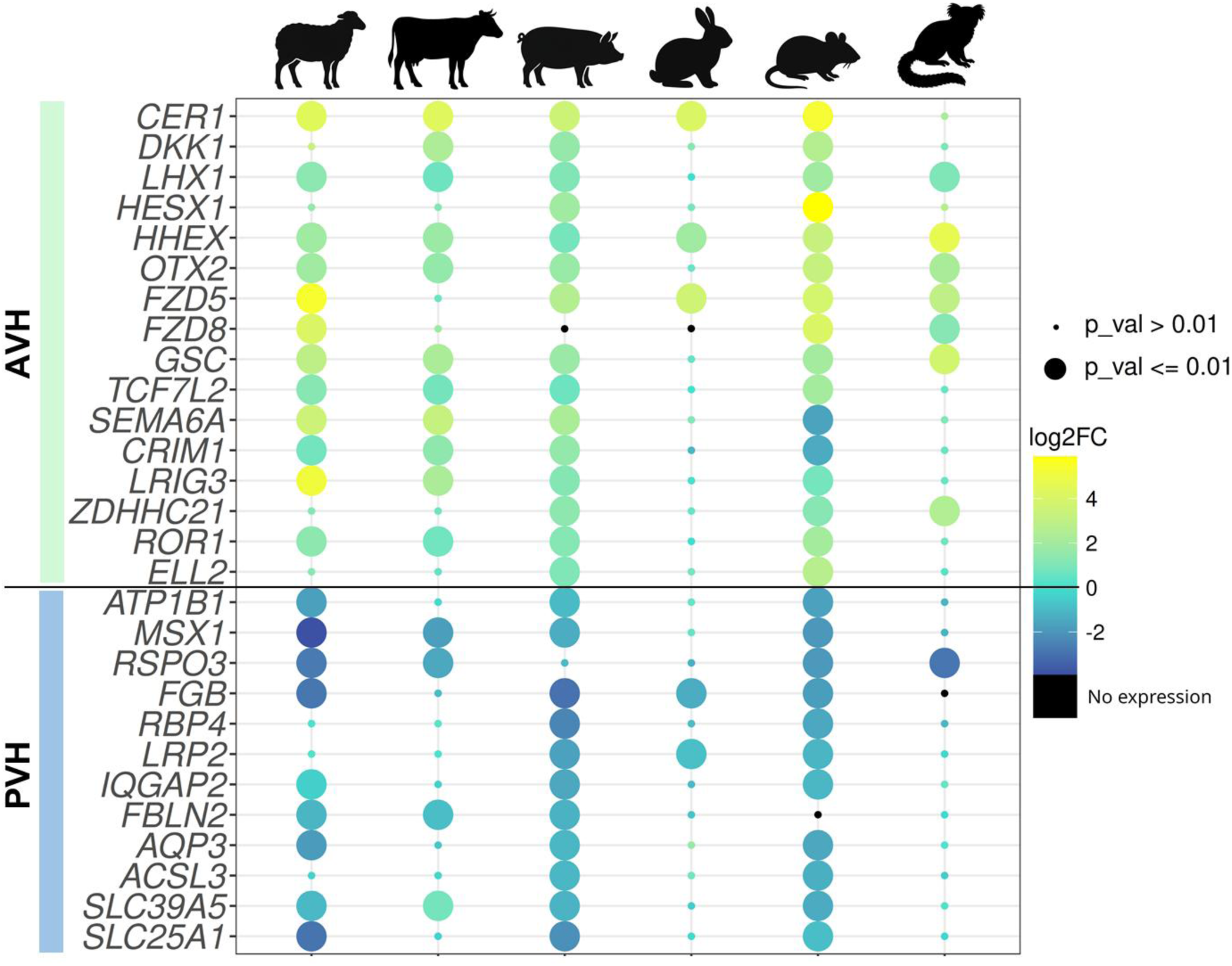
Pan-species anterior and posterior visceral hypoblast/endoderm markers. Bubble plot showing the log2 fold-change in gene expression between AVH and PVH clusters in early-PS embryos from sheep, cow, pig, rabbit, mouse, and marmoset. Bubble size indicates the p-value.

### Signalling pathways operating during symmetry breaking across species

To dissect inter-lineage communication, we applied CellChat ^45^ to pre-PS and early-PS stages across the six species (**Fig. 6a-l, Supplementary Fig. 5-16**). BMP, WNT, and FGF emerged as major signalling pathways mediating symmetry breaking in all species, consistent with previous findings ^46^.

**Figure 6.**
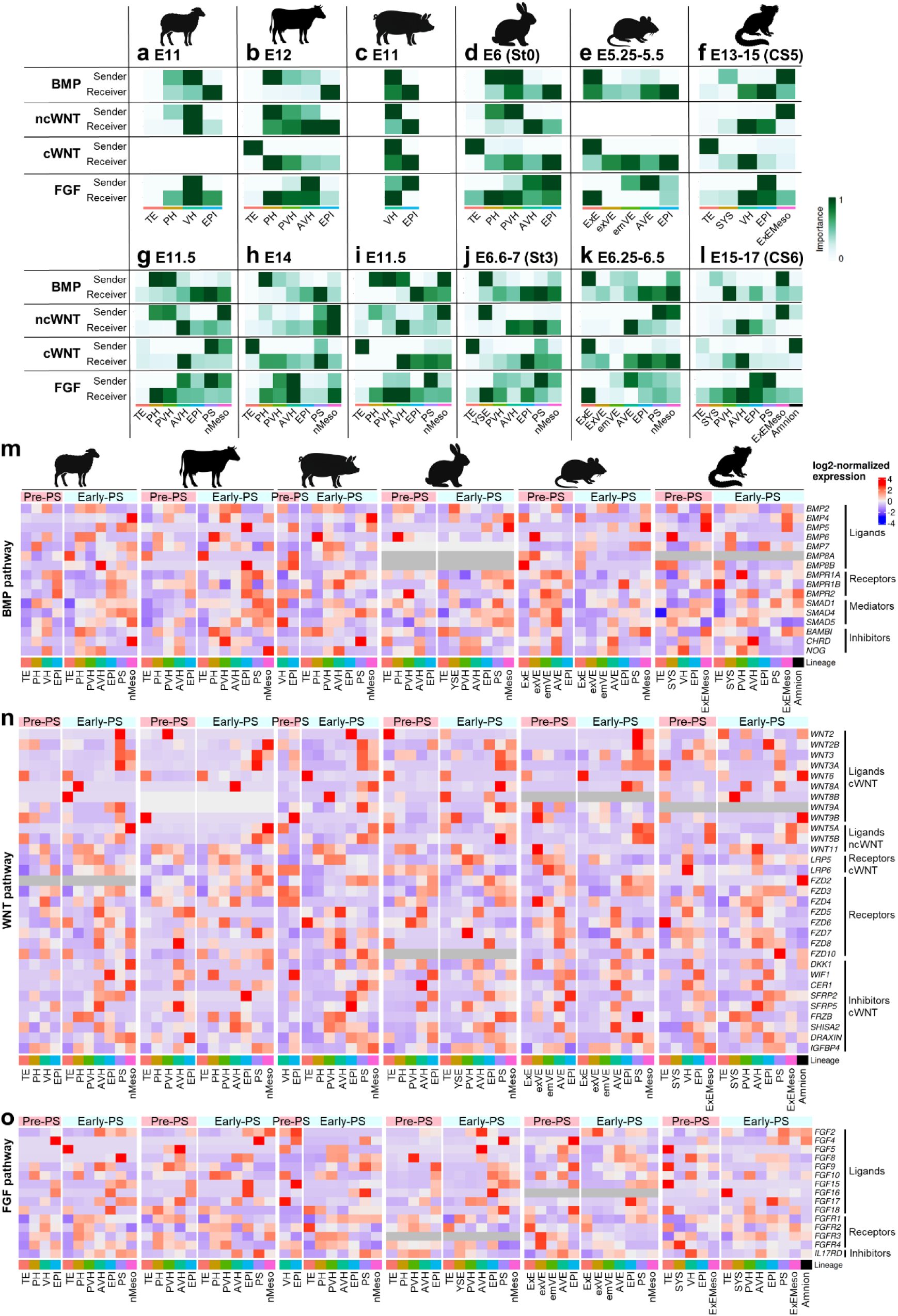
Cross-species comparison of signalling pathways mediating interlineage communication. a-l) Heatmaps showing the relative importance of the main signalling pathways mediating communication between all lineages in pre-PS (a-f) and early-PS (g-l) sheep, cow, pig, rabbit, mouse, and marmoset embryos. m-o) Heatmaps showing log2 normalized expression of ligands, receptors, mediators and inhibitors of m) BMP, n) WNT, and o) FGF pathways in sheep, cow, pig, rabbit, mouse, and marmoset pre-PS and early-PS embryos. AVH: anterior visceral hypoblast; EPI: epiblast; emVE: embryonic visceral endoderm; ExEct: extraembryonic ectoderm; ExEMeso: extraembryonic mesoderm; exVE: extraembryonic visceral endoderm; nMeso: nascent mesoderm; PH: parietal hypoblast; PS: primitive streak; PVH: posterior visceral hypoblast; SYS: secondary yolk sac; TE: trophectoderm; YSE: yolk sac endoderm.

In mouse embryos (**Fig. 6e and k, Supplementary Fig. 9 and 15**), TE-derived ExE was the primary BMP source, expressing *BMP4* from pre-PS stages to induce DVE/AVE formation ^30,31^ (**Fig. 4f**).

Conversely, in sheep (**Fig. 6a and g, Supplementary Fig. 5 and 11**), cow (**Fig. 6b and h, Supplementary Fig. 6 and 12**), pig (**Fig. 6c and i, Supplementary Fig. 7 and 13**), rabbit (**Fig. 6d and j, Supplementary Fig. 8 and 14**), and marmoset (**Fig. 6f and l, Supplementary Fig. 10 and 16**) embryos, the TE did not contribute to inter-lineage signalling, and *BMP4* was absent at the pre-PS stage in sheep and pigs or expressed at a very low level in cow and rabbit hypoblast and in marmoset ExEMeso. Instead of the TE, the hypoblast acted as the main BMP sender across non-rodent species, with BMP receptors expression enriched in EPI and PS (**Fig. 6m**). Across all species, *BMP2* was the predominant ligand expressed by hypoblast/endoderm (plus YSE in rabbit and SYS in marmoset), targeting the EPI at pre-PS and the PS at early-PS stages. Similar *BMP2* patterns have been reported in macaque and human embryos ^10^, suggesting that BMP2 may play a similar role in flat-disc species than BMP4 in mice. Several observations support this hypothesis: 1) BMP4 and BMP2 act via similar receptor complexes and show 92 % amino acid identity ^47,48^, 2) some *Bmp4* mutant mouse embryos progress beyond gastrulation ^47^ and form extraembryonic mesoderm progenitors, potentially rescued by embryonic *Bmp2* ^49^, 3) approximately half of *Elf5* mutant mouse embryos, lacking ExE and *Bmp4*, can develop ectopic T-positive mesoderm ^50^, and 4) *BMP2* has been implicated in PGC specification in mouse, possibly through additive effects of endoderm-derived *BMP2* and ExE-derived *BMP4*. In contrast, exposure to BMP2/BMP6 failed to induce *TBXT* in human ESCs ^10^, and thereby further functional studies are required in non-rodent embryos.

*BMP6* also played a conserved signalling role in ungulates and primates, secreted from PH to EPI (pre-PS) and to EPI, PS, and nMeso (early-PS) in sheep, cow, and pig, and from SYS to EPI (pre-PS) and EPI-derived structures (early-PS) in marmoset. *BMP6* enrichment has likewise been observed in peri-implantation human hypoblast ^10^. By early-PS, *BMP4* secretion by nMeso/ExEMeso was observed in all species, and by the amnion in marmoset. BMP inhibitors *BAMBI, CHRG* and *NOG* were expressed from the VH in all species, with stronger expression anteriorly, consistent with their role in maintaining pluripotency in the anterior EPI ^16^ (**Fig. 6m, Supplementary Fig. 5-16**).

Canonical WNT signalling was minimal at pre-PS in all species except marmoset, where ExEMeso expressed *WNT3/3A* (**Fig. 6a-f, Supplementary Fig. 10**). By early-PS, WNT activity increased in the posterior ED in all species, with *WNT3/3A* secreted by PS and nMeso (ExEMeso in marmoset) (**Fig. 6g-l, n**). Unlike mouse, where BMP4 induces *Wnt3* in the posterior epiblast to initiate PS formation ^47,51^, *BMP4* expression in non-rodent species was concomitant with, but not preceding AVH formation and WNT activation in the PS and nMeso. Thus, if BMP signalling is required for WNT induction in non-rodents, it may be mediated by VH-expressed *BMP2* or by PH-expressed *BMP6*, as previously discussed. WNT inhibitors such as *DKK1, CER1,* and *SHISA2* were expressed in AVH/AVE, while *WIF1* and *SFRP2/5,* among others, were expressed in the anterior epiblast to prevent differentiation and maintain pluripotency (**Fig. 6n**).

Non-canonical WNT ligands *WNT11*, *WNT5A/5B* were expressed in PS and nMeso (ExEMeso in marmoset), respectively (**Fig. 6n, Supplementary Fig. 5-16**). These genes have been implicated in EMT during A-P axis elongation in mouse ^52^ and may similarly regulate EMT during epiblast-to-mesoderm differentiation and migration.

FGF signalling also displayed species-specific differences. In mouse, *Fgf8* is the only FGF ligand required for gastrulation ^53^, but it was not detected during EPI-to-nMeso transition in the human gastrula ^18^ at later stages than those analysed in our study. In contrast, we detected *FGF8* and *FGF2* expression in VH, EPI, and PS, as well *FGF4* expression in EPI and PS across all species. However, the main FGF ligands in ungulates and lagomorphs were *FGF15* and *FGF16*, predominantly secreted by EPI-derived populations, and *FGF17*, mainly secreted by VH-derived cells (**Fig. 6o**). According to CellChat, *FGF15* and *FGF17* were key mediators of inter-lineage communication in ungulates and rabbit, but not in mouse and marmoset (**Supplementary Fig. 5-16**).

### NODAL is essential for epiblast and anterior visceral hypoblast development

Our pan-species analyses revealed conserved NODAL signalling dynamics during symmetry breaking across species, consistently with previous reports ^9,15,16,34,54^. *NODAL* was expressed in VH at both pre-PS and early-PS stages across all species, targeting the EPI at pre-PS and its derivatives (PS and ExEMeso) at early-PS. NODAL activators *FURIN* and *PCSK6 (PACE4)* were mainly expressed in extraembryonic lineages, while the NODAL co-receptor *CRIPTO* showed specific expression in the EPI and its derived lineages, consistent with observations in mouse post-implantation embryos ^54^. The NODAL inhibitors *CER1* and *LEFTY1/2* were expressed from the AVH/AVE across species (**Fig. 7a, Supplementary Fig. 5-16**), protecting the anterior EPI from mesoderm-inductive signals ^55^.

**Figure 7.**
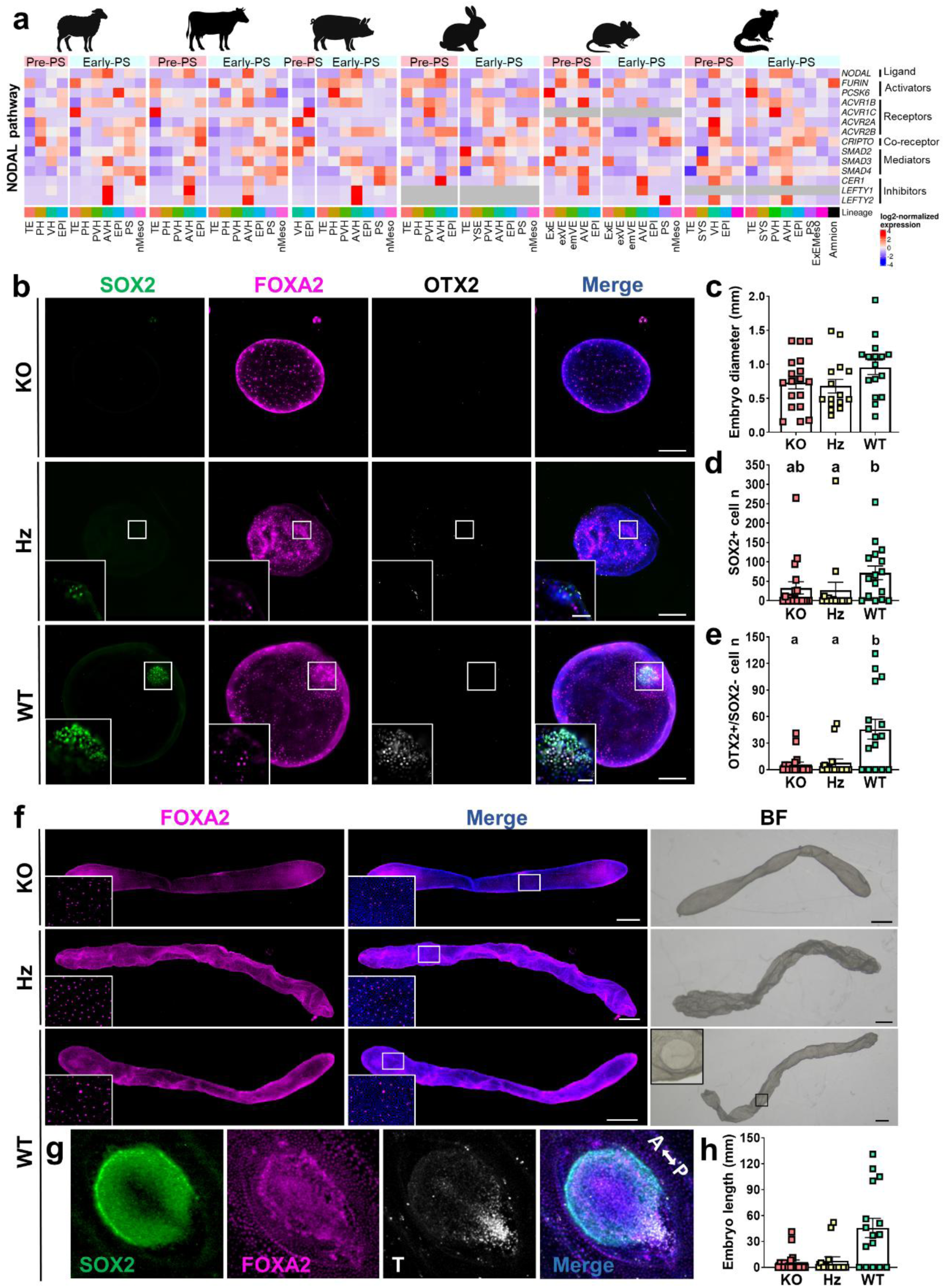
*NODAL* ablation impairs epiblast and anterior visceral hypoblast (AVH) survival in sheep embryos. a) Heatmap showing log2 normalized expression of ligand, activators, receptors, co-receptor, mediators and inhibitors of NODAL pathway in sheep, cow, pig, rabbit, mouse, and marmoset pre-PS and early-PS embryos. b) Representative immunofluorescence images of *NODAL* knock-out (KO), heterozygous (Hz) and wild-type (WT) E12 whole embryos and ED magnifications, stained for SOX2 (green; epiblast), FOXA2 (magenta; hypoblast) and OTX2 (white; epiblast and anterior visceral hypoblast). Nuclei were counterstained with DAPI (merge). White boxes in whole embryos images indicate magnified regions in the panels below. Scale bars: 200 µm for whole embryos and 50 µm for ED magnifications. c) Scatter plot showing embryo diameter (mean ± s.e.m.) of KO (n = 18), Hz (n = 15), and WT (n = 16) E12 embryos. The number of embryos analysed in each group is shown below columns. No statistically significant differences were detected (P > 0.05; one-way ANOVA). d-e) Scatter plots showing the number of SOX2+ epiblast cells and OTX2+/SOX2- anterior visceral hypoblast cells in KO (n = 18), Hz (n = 15), and WT (n = 16) E12 embryos (mean ± s.e.m.). The number of analysed embryos is shown in b). Different letters above columns indicate statistically significant differences (P ≤ 0.05; non-parametric Kruskal-Wallis test). f) Representative bright field (BF) and immunofluorescence images of *NODAL* KO, Hz and WT E14 elongated conceptuses stained for FOXA2 (magenta; hypoblast). Nuclei were counterstained with DAPI (merge). White boxes in whole embryos indicate magnified regions in the bottom left of the images, showing hypoblast migration along the inner TE surface. Scale bars: 1 mm. g) Representative immunofluorescence images of a WT E14 ED stained for SOX2 (green; epiblast), FOXA2 (magenta; hypoblast), and T (BRACHYURY, white; mesoderm). Nuclei were counterstained with DAPI (merge). Scale bar: 500 µm. h) Scatter plot showing embryo length of KO (n = 16), Hz (n = 2), and WT (n = 9) E14 embryos. The number of embryos analysed in each group is shown in each column. No statistically significant differences were detected (P > 0.05; non-parametric Kruskal-Wallis test).

However, functional studies outside rodents have so far been restricted to inhibitor treatments applied only up to the blastocyst stage ^10,56^. We therefore sought to investigate the role of NODAL during ED formation and symmetry breaking by generating *NODAL* knock-out (KO) sheep embryos using the cytosine base editor BE3 ^57^ to introduce premature stop codons near the start of the coding region (**Supplementary Fig. 17a**). *NODAL* expression in primates and ungulates starts in epiblast cells from the blastocyst stage ^16,38,56,58,59^. At day (D) 8, *in vitro* development to the blastocyst stage was comparable between sheep embryos microinjected with BE3 mRNA and sgRNA against *NODAL* (BE+G group; partially containing KO embryos) and control embryos injected with BE3 alone (BE group; WT embryos only) (**Table 1**). Among 21 genotyped blastocysts, 12 were classified as KO, corresponding to a direct KO efficiency of 57.14%. KO blastocysts displayed normal morphology, and lineage analysis revealed no significant differences among KO, heterozygous (Hz), and WT blastocysts in the number of SOX2+ epiblast, SOX17+ hypoblast, CDX2+ TE, or total cells (**Supplementary Fig. 17b, c**). These results indicate that NODAL is not required for blastocyst formation or early epiblast specification, consistent with recent findings in human embryos showing that NODAL inhibition from D5 to D7 blastocysts does not alter the expression of pluripotency factors ^56^.

**Table 1.** Total number of embryos and developmental rates in each experimental group at day (D) 8 and 12 of development *in vitro*, and at embryonic day (E) 12 and 14 of development *in vivo*.

| Day of development | Experimental replicates | Microinjected oocytes, n | Lysed oocytes (%) | Zygotes, n | Cleaved embryos (%) | Blastocysts at D7 (*in vivo*) / 8 (*in vitro*) (%) | D12 survival rates from D6/7 (%) |
| --- | --- | --- | --- | --- | --- | --- | --- |
| **D8 *in vitro*** | 2 |  |  |  |  |  | - |
| BE+G |  | 207 | 4 (1.93) | 203 | 182 (89.66) | 78 (38.42) |  |
| BE |  | 172 | 9 (5.23) | 163 | 150 (92.02) | 72 (44.17) |  |
| **D12 *in vitro*** | 3 |  |  |  |  |  |  |
| BE+G |  | 387 | 8 (2.06) | 379 | 338 (89.18) | 175 (51.77) | 136/175 (77.71) |
| BE |  | 216 | 11 (5.09) | 205 | 189 (92.19) | 105 (55.55) | 83/105 (79.05) |
| **E12 *in vivo*** | 2 |  |  |  |  |  | - |
| BE+G |  | 182 | 18 (9.89) | 164 | 137 (83.54) | 40 (24.39) |  |
| BE |  | 132 | 21 (15.91) | 111 | 96 (86.49) | 37 (33.33) |  |
| **E14 *in vivo*** | 1 |  |  |  |  |  | - |
| BE+G |  | 150 | 5 (3.33) | 145 | 127 (87.59) | 64 (44.14) |  |
| BE |  | 40 | 2 (5.00) | 38 | 32 (84.21) | 17 (44.74) |  |

No statistically significant differences were observed in developmental rates between the group microinjected with *NODAL* gRNA and BE3 mRNA (BE+G group, partially composed by KO embryos) and the group microinjected with BE3 only as a control (containing only WT embryos; Chi-square test; P>0.05). Cleavage and blastocyst rates are calculated from zygote numbers. Embryo survival at D12 is calculated from blastocyst placed in post-hatching *in vitro* culture.

We next examined development beyond the blastocyst stage using an extended *in vitro* culture system allowing ED development ^22^. Embryo survival from blastocyst to D12 was similar between *NODAL*-targeted (BE+G) and control (BE) groups (**Table 1**). Among 136 embryos analysed, 70 were KO (direct KO generation efficiency of 51.47%). EEMs development was unaffected by *NODAL* loss, as embryo growth and the extent of hypoblast migration along the inner TE surface were comparable among KO, Hz, and WT embryos. However, the number of SOX2+ and NANOG+ epiblast cells, as well as the number of AVH cells (OTX2+/SOX2-) were significantly lower in KO compared to WT embryos (**Supplementary Fig. 18**).

To validate these findings, and to assess later developmental stages, D7 blastocysts from both *NODAL*-targeted and control groups were transferred to recipient ewes. Embryo growth and hypoblast migration remained unaffected when conceptuses were recovered after five (E12) or seven (E14) days. However, epiblast development was significantly impaired in KO and Hz embryos compared to WT, validating the results found *in vitro*. At E12, most KO and Hz E12 embryos lacked an ED, and some displayed degenerating EDs with sharply reduced SOX2+ epiblast cell numbers. The AVH was also compromised, as both the proportion of embryos showing surviving AVH cells (OTX2+/SOX2-) and the number of AVH cells were significantly lower in KO and Hz compared to WT (**Fig. 6 b-e,d, Table 2**). This aligns with findings in human peri-implantation embryos treated with a NODAL inhibitor between D7 and D9, where the number of CER1+ AVH cells decreased ^10^. At E14, all KO and Hz embryos lacked an ED, whereas 66.67% of WT embryos showed gastrulating EDs (**Fig. 6 f-h, Table 2**). Similarly, *Nodal* mutant mouse embryos arrest before gastrulation, fail to express *Otx2* in the visceral endoderm ^60^, cannot induce DVE/AVE specification, and do not maintain epiblast pluripotency ^54^.

**Table 2.** Hypoblast and epiblast development in *NODAL* knockout (KO), heterozygous (HZ) and wild-type (WT) conceptuses recovered at embryonic days (E) 12 and 14.

| Embryonic day | Genotype | Complete hypoblast migration (%) | Visceral hypoblast survival (OTX2+/SOX2-; %) | Epiblast survival (SOX2+; %) | ED formation (%) |
| --- | --- | --- | --- | --- | --- |
| 12 | KO | 10/18 (55.55) | 7/18 (38.89%)<sup>a</sup> | 9/18 (50)<sup>ab</sup> | 6/18 (33.33)<sup>a</sup> |
|  | HZ | 5/15 (33.33) | 4/15 (26.67%)<sup>a</sup> | 5/15 (33.33)<sup>a</sup> | 3/15 (20)<sup>a</sup> |
|  | WT | 7/16 (43.75) | 12/16 (75%)<sup>b</sup> | 13/16 (81.25)<sup>b</sup> | 12/16 (75)<sup>b</sup> |
| 14 | KO | 14/16 (87.5) | - | 0/16 (0)<sup>a</sup> | 0/16 (0)<sup>a</sup> |
|  | HZ | 2/2 (100) | - | 0/2 (0)<sup>ab</sup> | 0/2 (0)<sup>ab</sup> |
|  | WT | 9/9 (100) | - | 6/9 (66.67)<sup>b</sup> | 6/9 (66.67)<sup>b</sup> |

KO: knock-out; HZ: heterozygous; WT: wild-type; ED: embryonic disc. Different letters (a, b) indicate statistically significant differences (Chi-square test; $P \leq 0.05$ ).

Together, these findings demonstrate that while NODAL is dispensable for early epiblast formation at the blastocyst stage, it is essential for epiblast maintenance and AVH development during ED formation and symmetry breaking in sheep.

## Discussion

Although the majority of pregnancy failures in primates and ungulates occur during peri-implantation, the molecular regulation of this developmental window remains poorly understood. A defining feature of this period across these species is the formation of a flat ED that must undergo symmetry breaking and gastrulation. By generating a single-cell transcriptomic atlas of *in vivo*-derived sheep embryos from pre-PS to late-PS stages, we provide a high-resolution framework of the onset of symmetry breaking and gastrulation in a mammalian model forming an ED. Our analyses reveal the coordinated timing of lineage segregation and uncover dynamic differences in lineage proliferation, including rapid expansion of mesoderm in contrast to the slower proliferation of the endoderm, consistent with previous observations in other species ^15^.

To place sheep development in a broader evolutionary context, we assembled a cross-species atlas of symmetry breaking across six mammals. This comparative approach identified conserved AVH and PVH markers and revealed the molecular pathways underlying anterior–posterior axis establishment. Despite substantial interspecies differences in embryo morphology, developmental timing, and the contribution of extraembryonic lineages, the major signalling pathways that orchestrate symmetry breaking (NODAL, BMP, WNT, and FGF) were conserved across mammals, with mediators of WNT and NODAL signalling being particularly well conserved. By combining transcriptomics with loss-of-function studies, we demonstrate that NODAL is dispensable for early epiblast specification in sheep but essential for maintaining epiblast and AVH survival during symmetry breaking. These findings support a pan-species model in which NODAL sustains epiblast pluripotency, while anteriorly expressed WNT inhibitors protect this domain from mesoderm induction. Conversely, conserved posterior WNT ligands promote mesoderm differentiation, highlighting a shared regulatory logic across mammals ^14–16^.

FGF signalling also supports epiblast pluripotency across species, although our dataset revealed notable divergence in ligand expression, with ungulates and lagomorphs relying on different FGF ligands than primates and rodents. BMP signalling displayed the most pronounced interspecies differences. In mice, *BMP4* is expressed by the TE and acts as a major inducer of gastrulation, whereas in sheep, cow, pig, and rabbit the hypoblast constitutes the primary BMP source, and instead of BMP4, other ligands such as BMP2 and BMP6 appear to play dominant roles. In primates, in addition to the hypoblast, early ExEMeso and the amnion also likely contribute BMP signals 16,61,62.

These species-specific variations have important implications for human developmental biology. Key embryo structures, such as the AVH, early ExEMeso, and amnion, do not differentiate robustly in current extended human embryo culture systems or in stem cell–based embryo models, which are heavily influenced by mouse-based developmental paradigms ^12,20^. In our analyses, integration of human embryos into the cross-species atlas was hindered by the lack of equivalent *in vivo* datasets and by insufficient representation of key lineages in available *in vitro* datasets ^29,63^. These challenges underscore the need for alternative models, such as sheep and other ungulates or rabbits, which develop a flat ED and are highly accessible due to their relevance in agriculture. These species provide a tractable and physiologically relevant platform for decoding the molecular networks governing early human development.

Limitations of our study include the incomplete availability of equivalent developmental stages across species, the absence of a comprehensive *in vivo* human embryo dataset for direct comparison, and variability in the quality of genome assemblies, transcriptomic depth, and gene annotations among species, which may influence data interpretation.

Overall, our work presents a detailed molecular atlas of symmetry breaking in a mammal developing a flat ED and highlights both universal principles and species-specific mechanisms shaping mammalian symmetry breaking and gastrulation. By integrating comparative single-cell transcriptomics, CRISPR-based genome editing, extended embryo culture, and lineage tracing, our study underscores the utility of the sheep as a powerful and accessible model species for developmental biology. These findings provide a foundation for improving human stem cell-based embryo models and for advancing our understanding of how the body plan is established across mammalian embryos that develop a flat embryonic disc.

## Methods

### Ethical statement

All experimental procedures were approved by INIA Animal Care Committee and Madrid Region Authorities (PROEX 313.0/21) in agreement with European legislation.

### In vivo recovery of ovine embryos

*In vivo* embryos were obtained from 3 years old Merino ewes superovulated following the protocol described in **Supplementary Fig. 18**, which involved flugestone acetate (0.35 g, CIDR® OVIS, Zoetis), PGF2α analog (Cloprostenol, Estrumate®, MSD), luteinizing and follicle-stimulating hormone (Pluset, Callier). Ewes were mated (day 0 PM and day 1 AM) and slaughtered at days 11, 11.5, 12.5 and 13.5 post-coitus. Embryos were recovered by uterine flushing with warmed recovery medium (Euroflush, IVM Technologies).

### In vitro production of NODAL KO ovine embryos

Ovine *NODAL* KO embryos were generated by introducing a stop codon at the first exon using the cytosine base editor BE3 ^64^. A single guide RNA (sgRNA) was designed to convert two TGG codons into TAG/TGA/TAA stop codons and thereby disrupt protein formation. The sgRNA (CTTGCACGCC**TGGTGG**GCTC) was produced using the Guide-it sgRNA *in vitro* Transcription kit® (Takara Bio, USA) using a primer containing a T7 promoter and the sgRNA sequence (**Supplementary Table 1**). Capped polyadenylated BE3-encoding mRNA was produced by *in vitro* transcription using the mMESSAGE mMACHINE T7 ULTRA kit (Life Technologies) and the plasmid pCMV-BE3 (Addgene #73021) linearized with BbsI. mRNA was purified using the MEGAClear kit (Life Technologies).

Cumulus oocyte complexes (COCs) were aspirated from 2 to 8 mm diameter follicles of ovine ovaries collected at a local slaughterhouse using a 21 G needle connected to an aspiration pump (VMAR 5100, Cook) adjusted to 25 mmHg. COCs were collected in a 50 ml tube containing Euroflush medium (IMV Technologies, France). COCs with compact cumulus and homogeneous cytoplasm were selected and matured for 22 h in TCM-199 supplemented with 40 µg/ml gentamicin sulphate, 10% (v/v) fetal bovine serum (FBS) and 10 ng/ml epidermal growth factor at 38.5 °C under an atmosphere of 5% CO2 in air with maximum humidity. Then, matured oocytes were denuded by vortexing for 3 min in 1 ml of 300 µg/ml hyaluronidase in phosphate-buffered saline (PBS), and intra-cytoplasmic microinjection was performed with a spike-end micropipette (5 µm internal diameter; BioMedical Instruments, Germany) connected to a manual hydraulic air microinjector (CellTram Air, Eppendorf, Germany) under a Nikon Eclipse TE300 microscope, as previously described ^65^.

Oocytes were randomly allocated into two groups: i) *NODAL*-targeted group, microinjected with 200 ng/µl BE3 mRNA and 100 ng/µl *NODAL* sgRNA (approximately 2/3 of the oocytes per replicate), and ii) non-targeted group (control), microinjected with 200 ng/µl BE3 mRNA (1/3 of the oocytes per replicate). The control group served as a microinjection control in which no genome edition is expected.

Immediately after microinjection (24 h post-*in vitro* maturation), oocytes were fertilized in groups of approximately 25 with frozen-thawed Bovi-Pure® (Nidacon, Sweden)-separated ram sperm at a final concentration of 2 × 10^6^ spermatozoa/ml. Gametes were co-incubated in 50 µl droplets of IVF medium (Stroebech media, Denmark) supplemented with 10% (v/v) heat-inactivated oestrous sheep serum and 50 IU/ml heparin, covered by mineral oil, at 38.5 °C in an atmosphere of 5% CO2 with maximum humidity. Semen from the same ram was used for all experimental replicates to avoid a potential confounding effect of individual ram variability on embryo development.

At 18 h post-insemination, presumptive zygotes were denuded by manual pipetting and cultured in groups of approximately 25 in 50 µl droplets of IVC medium (Stroebech media, Denmark), covered by mineral oil, at 38.5 °C under an atmosphere of 5% CO2, 5% O2 and 90% N2 with maximum humidity. Cleavage was assessed at 48 h and blastocyst rates were recorded at day (D) 8.

### Post-hatching development system

At D6 and D7 post-fertilisation, blastocysts were transferred to Nunclon Sphera low-attachment dishes (Thermo Scientific, Denmark) and cultured in N2B27 medium (1:1 Neurobasal and DMEM/F12 medium supplemented with penicillin/streptomycin, 2 mM L-glutamine and N2 and B27 supplements (Thermo Fisher Scientific) ^22,66^ at 38.5 °C under an atmosphere of 5% CO2, 5% O2 and 90% N2 with maximum humidity. Half of the culture medium was replaced every other day until D12, when pictures were taken and embryos were fixed.

### In vivo development of NODAL KO conceptuses

All experimental procedures were approved by INIA Animal Care Committee and Madrid Region Authorities (PROEX 059.2/22) in compliance with European legislation.

D7 blastocysts from the *NODAL*-targeted (partially composed of KO embryos) and control (composed by wild-type [WT] embryos only) groups were transferred to three recipient ewes (**Supplementary Table 2**). Oestrus synchronization was achieved by inserting a 0.35 g progesterone-releasing device (CIDR-Ovis, Zoetis, USA). Cloprostenol (100 µg, Estrumate®, MSD) was administered 11 days after CIDR insertion, and PMSG (450 IU, Sincropart PMSG, CEVA, France) was administered on the day of CIDR removal (12 days post-insertion). Embryo transfer took place 8 days post-CIDR removal. The presence of corpora lutea was assessed by abdominal laparoscopy, and the cranial section of the uterine horns was exteriorized to introduce the embryos. After 5 or 7 days, ewes were slaughtered and embryonic day (E) 12 or 14 embryos (**Supplementary Table 2**) were recovered by uterine flushing with Euroflush® medium (IMV Technologies, France). Embryos were photographed to measure size and then fixed for further analyses.

### Immunofluorescence and lineage development analysis

Embryos were fixed in 4 % paraformaldehyde (PFA) for 15 minutes at room temperature (RT), washed in PBS supplemented with 1 % BSA, permeabilized in 1 % Triton X-100 in PBS for 15 min at RT, and blocked in 10 % Donkey Serum with 0.02 % Tween 20 in PBS for 1 h at RT. Embryos were then incubated overnight at 4 °C with primary antibodies (**Supplementary Table 3**). After four washes in PBS-1% BSA, embryos were incubated in blocking solution containing DAPI and the appropriate Alexa-conjugated secondary antibodies (**Supplementary Table 3**) for 1 h at RT, followed by four additional washes in PBS-1 % BSA. Finally, embryos were mounted on PBS-1% bovine serum albumin (BSA) microdrops made by drawing circles with a PAP pen (Kisker Biotech GmbH) on a coverslide as previously described (Ramos-Ibeas, Lamas-Toranzo et al. 2020). Microdrops were covered by an Invitrogen^TM^ CoverWell^TM^ incubation chamber gasket (Fisher Scientific) to prevent crushing. Imaging was performed using a Zeiss Axio Observer microscope equipped with ApoTome.2 structured illumination (Zeiss).

Following image acquisition, embryo diameter was measured to assess size, and lineage development was analysed. Total, SOX2+, SOX17+ and CDX2+ cell numbers in D8 and D12 *in vitro* embryos, as well as SOX2+ and OTX2+ cell numbers in E12 and E14 *in vivo* conceptuses, were manually counted using ZEN 2.6 software (Zeiss). In D12 *in vitro* embryos, hypoblast migration was quantified as the extension of the SOX17+ hypoblast layer along the inner surface of the CDX2+ trophectoderm. Each spherical embryo was bisected along the Z axis, and orthogonal projections were generated. Total area and the area covered by SOX17+ hypoblast cells were measured in each projection, and hypoblast migration was expressed as the percentage of hypoblast-covered area relative to the total, averaged across both projections. Epiblast survival was defined by the presence of SOX2+ epiblast cells in the embryo, and embryonic disc (ED) formation was identified as a compact cluster of ≥ 30 SOX2+ cells.

### Embryo genotyping by Sanger sequencing

Following immunofluorescence, embryos (whole D8 or D12 *in vitro*; whole E12 *in vivo* embryos or fragments of E14 *in vivo* embryos) were placed at the bottom of 0.2 ml PCR tubes and stored at −20 °C until analysis. Samples were digested with 8 µl (whole embryos) or 10 µl (E14 fragment) of Arcturus Picopure® DNA extraction solution (ThermoFisher Scientific), incubated at 65 °C for 1 h, and heat-inactivated at 95 °C for 10 min. Lysates were subsequently used to amplify the BE3 target region by PCR with primers listed in **Supplementary Table 1**. Reactions contained 2 µl of lysate in 25 µl PCR reaction (GoTaq Flexi, Promega). PCR conditions were: 96 °C for 2 min; ×40 (96 °C for 20 s, 64 °C for 30 s, 72 °C for 30 s); 72 °C for 5 min; hold at 8 °C. PCR products were purified by FavorPrep PCR purification kit (Favorgen) and Sanger sequenced. Embryos were classified as WT (containing no mutated alleles, i.e., all embryos in the control group), heterozygotes (containing WT and KO alleles) and KO (harboring only alleles containing the stop codon).

### In vivo embryo digestion and cells fixation

Embryos were digested and fixed right after flushing to avoid RNA degradation. To mitigate the overrepresentation of trophectoderm and hypoblast cells in elongated conceptuses, embryos from E12.5 onwards were dissected with scalpels and forceps to remove most of these cells, and only the embryonic disc and surrounding extraembryonic membranes (hypoblast and trophectoderm) were isolated for subsequent cell digestion. Embryos were chopped and dissociated into single cells by incubation in TrypLE in 4-well dishes in an orbital shaker at 150 rpm, for 20 min at 38.5 °C. During this time, embryos were pipetted every 3 to 5 minutes and finally washed with ice-cold Euroflush medium (IVM Technologies) to quench TrypLE activity. Cells were filtered through Flowmi 40 µm cell strainers, collected in BSA-coated Falcon tubes and centrifuged at 300 G at 4 °C for 8 min. Then, cells were fixed using the Evercode^TM^ Cell Fixation kit v2 (ECF2101, Parse Biosciences) following the manufacturer instructions. Briefly, cells were suspended in Cell Prefixation Buffer, stained with Trypan Blue to assess cell viability and counted using a haemocytometer. Then, Cell Fixation Solution was added and cells were incubated on ice for 10 minutes. Next, Cell Permeabilization Solution was added, and after 3 min incubation on ice, Cell Neutralization Buffer was added to block the reaction. Then, cells were centrifuged at 300 G at 4 °C for 8 min, suspended in Cell Buffer and frozen in a Mr. Frosty^TM^ freezing container at −80 °C until libraries preparation.

### Preparation of scRNAseq libraries and sequencing

Single-cell libraries were constructed from 9 *in vivo* embryo samples collected at different stages (**Supplementary Data 1**) using the Evercode^TM^ WT Mini v2 kit (Parse Biosciences) according to the manufacturer’s protocol. This plate-based combinatorial barcoding method allowed to generate two sublibraries with around 12,000 cells each.

cDNA sub-libraries were assessed using Qubit dsDNA HS kit (Thermo Fisher Scientific) and Agilent 2100 Bioanalyzer (Agilent Technologies) and sent to an external company (Azenta Life Sciences, Germany) to undergo 150 bp paired-end sequencing (1,750 × 10^6^ total reads) using the NovaSeq 6000 platform.

### Data availability and scRNAseq data preprocessing

The scRNAseq datasets generated during this study are available under GEO accession number: GSE320427. Data were processed with the Parse Biosciences pipeline (v1.3.1) using default settings. Samples were demultiplexed and sequencing reads were aligned to the ovine genome (ARS-UI_Ramb_v2.0, Ensembl release 113). Downstream analyses were performed using the R package Seurat (v5.3.0).

### Filtering of cells, integration, dimensionality reduction and clustering

Expression matrices were loaded with the *ReadParseBio* function, and Seurat objects were generated for each of the 9 samples, which were independently processed. Cells with more than 20,000-60,000 detected reads, fewer than 1,000-3,000 or more than 5,000-9,000 detected genes were excluded from analyses. Cells were not filtered based on mitochondrial transcript content, given the heterogeneity embryonic cell types; however, mitochondrial gene percentages were on average lower than 1% in all samples, except for E11 (∼2%). The gene-cell matrices were normalized and scaled (*NormalizeData* and *ScaleData* functions), and the 3,000 most variably expressed genes were identified using the *FindVariableFeatures* function. These genes were used to calculate the first 50 principal components (PCs) with *RunPCA* function for individual samples, and the first 100 PCs for the integrated samples.

Cells clustering was performed with *FindNeighbors* (20 dimensions of reduction for individual samples, 50 for the integrated object) and *FindClusters* functions (resolution adjusted between 0.2 and 0.8 according to the cell lineage complexity observed at each developmental stage). Clusters were visualized using *RunUMAP* function. Cluster identities were assigned based on the expression of established lineage-specific markers reported in previous single-cell transcriptomic studies across species (**Supplementary Data 2**). In addition, new cell markers were identified by differential expression analysis, applying the Wilcoxon rank-sum test to the integrated Seurat object. Genes were considered as differentially expressed (DEGs) and thus cluster-specific markers if they met the following criteria: logFC > 2, adjusted p value < 0.01, expression ≥ 30% of cells within the cluster, and expression in ≤ 30% of cells in all other clusters.

Cluster 6 co-expressed both TE and hypoblast markers (**Supplementary Fig. 2e**), suggesting a mixed or transitional identity, so it was excluded from subsequent analyses.

### Gene ontology and KEGG pathway analysis

Gene Ontology (GO) and KEGG pathway analyses of cluster-specific markers were performed with *clusterProfiler* (v 4.16.0). Briefly, gene symbols corresponding to cluster markers were converted to Entrez IDs using the *bitr* function and analyzed with *compareCluster* function under default parameters. GO terms and KEGG pathways with an adjusted p-value < 0.01 were considered significantly enriched.

### Inter-species comparisons

The transcriptomes of pre-primitive streak (PS) and early-PS embryos from other mammalian species were also analysed. Either *Seurat* or *h5ad* AnnData objects were obtained from publicly available datasets for pig ^15,38^, marmoset ^16^, rabbit ^14^, and mouse ^11,39–41^. Further information on methods for inter-species comparisons is provided in the Supplementary Information.

### Cell-cell communication analysis

Cell-cell signaling pathways in each species and selected developmental stages (pre-PS and early-PS) were analyzed using the R package *CellChat* (v2.1.2). The original *CellChat* ligand-receptor database was modified to incorporate *CRIPTO*/*TDGF1* as a co-receptor in the NODAL signalling pathway by duplicating *CFC1* entries and replacing the duplicated copy with *TDGF1*. After subsetting the data using *subsetData()* and identifiying overexpressed genes and ligand-receptor interactions in each lineage, gene expression values were smoothed, and inter-lineage communication probabilities and interaction strengths were inferred using *computeCommonProb().* The *triMean* method was applied for all species except rabbit, for which the less stringent *trunc*at*edMe*a*n* method was used to enhance interaction detection after an initial analysis with the *triMean* method failed to identify sufficient interactions. Plotting functions were customized to adjust font size, angle, color palette and displayed clusters, as coded in the GitHub script.

### Statistical analysis

Statistical analyses of *NODAL* KO embryos were performed using GraphPad Prism (GraphPad Software, San Diego, CA, USA), with statistical significance defined as P ≤ 0.05. Differences in developmental rates between the group microinjected with *NODAL* gRNA and BE3 mRNA (BE+G group, partially composed by KO embryos) and the group microinjected with BE3 alone (control group, containing only WT embryos) were assessed using the Chi-square test. Complete hypoblast migration rate, visceral hypoblast and epiblast survival, and ED formation rates in *in vivo*-developed embryos were also evaluated using the Chi-square test. Cell counts, embryo area, and the percentage of hypoblast migration were analysed using one-way ANOVA. When the DߣAgostino & Pearson normality test test failed, statistical differences were analyzed using the non-parametric Kruskal-Wallis test.

## Funding and acknowledgements

This work has been funded by the projects PID2021-122153NA-I00 and PID2024-155682NB-I00 from the Spanish Ministry of Science and Innovation to PRI, and StG-757886-ELONGAN from the European Research Council and ECQ2018-005184-P from the Spanish Ministry of Science and Innovation to PBA. LGB was funded by a JDC2022-049435-I Fellowship from the Spanish Ministry of Science and Innovation.

The authors want to acknowledge the staff of the slaughterhouse “Matadero Mondejano SL” for kindly providing sheep ovaries used in this study.

## Author contributions

L. G-B. performed bioinformatics analyses, advised on the project and wrote the paper; N. M-R. contributed to ovine *in vitro* embryo production, generated *NODAL* KO embryos, performed IF analyses and embryo genotyping; P. M. collaborated to ovine *in vitro* embryo production; A. T-D. and J. S.-M. were involved in embryo transfer experiments; L. S. and R. A. collaborated to bioinformatics analyses and advised on the project; P. B-A. performed *in vivo* embryo collection, dissection, and embryo transfers, and provided funding; P. R-I. designed and performed the experiments including embryo dissection and scRNA-Seq libraries generation, contributed to *in vitro* embryo production, analysed the results, provided funding, supervised the project and wrote the paper. All authors revised the manuscript.

## References

1 Tam, P. P. & Loebel, D. A. Gene function in mouse embryogenesis: get set for gastrulation. Nat Rev Genet 8, 368–381 (2007). 10.1038/nrg2084

2 Arnold, S. J. & Robertson, E. J. Making a commitment: cell lineage allocation and axis patterning in the early mouse embryo. Nat Rev Mol Cell Biol 10, 91–103 (2009). 10.1038/nrm2618

3 Perez-Gomez, A., Gonzalez-Brusi, L., Bermejo-Alvarez, P. & Ramos-Ibeas, P. Lineage Differentiation Markers as a Proxy for Embryo Viability in Farm Ungulates. Frontiers in Veterinary Science 8 (2021). 10.3389/fvets.2021.680539

4 Reese, S. T. et al. Pregnancy loss in beef cattle: A meta-analysis. Anim Reprod Sci 212, 106251 (2020). 10.1016/j.anireprosci.2019.106251

5 Bidarimath, M. & Tayade, C. Pregnancy and spontaneous fetal loss: A pig perspective. Mol Reprod Dev 84, 856–869 (2017). 10.1002/mrd.22847

6 Larsen, E. C., Christiansen, O. B., Kolte, A. M. & Macklon, N. New insights into mechanisms behind miscarriage. BMC Med 11, 154 (2013). 10.1186/1741-7015-11-154

7 Eakin, G. S. & Behringer, R. R. Diversity of germ layer and axis formation among mammals. Semin Cell Dev Biol 15, 619–629 (2004). 10.1016/j.semcdb.2004.04.008

8 Beddington, R. S. & Robertson, E. J. Axis development and early asymmetry in mammals. Cell 96, 195–209 (1999). 10.1016/s0092-8674(00)80560-7

9 van Leeuwen, J., Berg, D. K. & Pfeffer, P. L. Morphological and Gene Expression Changes in Cattle Embryos from Hatched Blastocyst to Early Gastrulation Stages after Transfer of In Vitro Produced Embryos. PLoS One 10, e0129787 (2015). 10.1371/journal.pone.0129787

10 Weatherbee, B. A. T. et al. Distinct pathways drive anterior hypoblast specification in the implanting human embryo. Nat Cell Biol 26, 353–365 (2024). 10.1038/s41556-024-01367-1

11 Pijuan-Sala, B. et al. A single-cell molecular map of mouse gastrulation and early organogenesis. Nature 566, 490–495 (2019). 10.1038/s41586-019-0933-9

12 Zhu, Q. et al. Decoding anterior-posterior axis emergence among mouse, monkey, and human embryos. Dev Cell 58, 63–79.e64 (2023). 10.1016/j.devcel.2022.12.004

13 Ton, M. N. et al. An atlas of rabbit development as a model for single-cell comparative genomics. Nat Cell Biol 25, 1061–1072 (2023). 10.1038/s41556-023-01174-0

14 Mayshar, Y. et al. Time-aligned hourglass gastrulation models in rabbit and mouse. Cell 186, 2610–2627.e2618 (2023). 10.1016/j.cell.2023.04.037

15 Simpson, L. et al. A single-cell atlas of pig gastrulation as a resource for comparative embryology. Nat Commun 15, 5210 (2024). 10.1038/s41467-024-49407-6

16 Bergmann, S. et al. Spatial profiling of early primate gastrulation in utero. Nature 609, 136–143 (2022). 10.1038/s41586-022-04953-1

17 Nakamura, T. et al. A developmental coordinate of pluripotency among mice, monkeys and humans. Nature 537, 57–62 (2016). 10.1038/nature19096

18 Tyser, R. C. V. et al. Single-cell transcriptomic characterization of a gastrulating human embryo. Nature 600, 285–289 (2021). 10.1038/s41586-021-04158-y

19 Xiao, Z. et al. 3D reconstruction of a gastrulating human embryo. Cell 187, 2855–2874.e2819 (2024). 10.1016/j.cell.2024.03.041

20 Mallapaty, S. Human embryo models are getting more realistic - raising ethical questions. Nature 633, 268–271 (2024). 10.1038/d41586-024-02915-3

21 Guillomot, M., Turbe, A., Hue, I. & Renard, J. P. Staging of ovine embryos and expression of the T-box genes Brachyury and Eomesodermin around gastrulation. Reproduction 127, 491–501 (2004). 10.1530/rep.1.00057

22 Ramos-Ibeas, P. et al. In vitro culture of ovine embryos up to early gastrulating stages. Development 149 (2022). 10.1242/dev.199743

23 Berg, D. K. et al. Trophectoderm lineage determination in cattle. Dev Cell 20, 244–255 (2011). 10.1016/j.devcel.2011.01.003 10.1016/j.devcel.2011.01.003.

24 Davenport, K. M. et al. Single-nuclei RNA sequencing (snRNA-seq) uncovers trophoblast cell types and lineages in the mature bovine placenta. Proc Natl Acad Sci U S A 120, e2221526120 (2023). 10.1073/pnas.2221526120

25 Johnson, G. A. et al. Cellular events during ovine implantation and impact for gestation. Anim Reprod 15, 843–855 (2018). 10.21451/1984-3143-AR2018-0014

26 Solloway, M. J. & Robertson, E. J. Early embryonic lethality in Bmp5;Bmp7 double mutant mice suggests functional redundancy within the 60A subgroup. Development 126, 1753–1768 (1999). 10.1242/dev.126.8.1753

27 Scialdone, A. et al. Resolving early mesoderm diversification through single-cell expression profiling. Nature 535, 289–293 (2016). 10.1038/nature18633

28 Saykali, B. et al. Distinct mesoderm migration phenotypes in extra-embryonic and embryonic regions of the early mouse embryo. Elife 8 (2019). 10.7554/eLife.42434

29 Molè, M. A. et al. A single cell characterisation of human embryogenesis identifies pluripotency transitions and putative anterior hypoblast centre. Nat Commun 12, 3679 (2021). 10.1038/s41467-021-23758-w

30 Rodriguez, T. A., Srinivas, S., Clements, M. P., Smith, J. C. & Beddington, R. S. Induction and migration of the anterior visceral endoderm is regulated by the extra-embryonic ectoderm. Development 132, 2513–2520 (2005). 10.1242/dev.01847

31 Richardson, L., Torres-Padilla, M. E. & Zernicka-Goetz, M. Regionalised signalling within the extraembryonic ectoderm regulates anterior visceral endoderm positioning in the mouse embryo. Mech Dev 123, 288–296 (2006). 10.1016/j.mod.2006.01.004

32 Takaoka, K., Yamamoto, M. & Hamada, H. Origin and role of distal visceral endoderm, a group of cells that determines anterior-posterior polarity of the mouse embryo. Nat Cell Biol 13, 743–752 (2011). 10.1038/ncb2251

33 Hoshino, H., Shioi, G. & Aizawa, S. AVE protein expression and visceral endoderm cell behavior during anterior-posterior axis formation in mouse embryos: Asymmetry in OTX2 and DKK1 expression. Dev Biol 402, 175–191 (2015). 10.1016/j.ydbio.2015.03.023

34 Yoshida, M. et al. Conserved and divergent expression patterns of markers of axial development in eutherian mammals. Dev Dyn 245, 67–86 (2016). 10.1002/dvdy.24352

35 Kemp, C. R. et al. Expression of Frizzled5, Frizzled7, and Frizzled10 during early mouse development and interactions with canonical Wnt signaling. Dev Dyn 236, 2011–2019 (2007). 10.1002/dvdy.21198

36 de Lau, W., Peng, W. C., Gros, P. & Clevers, H. The R-spondin/Lgr5/Rnf43 module: regulator of Wnt signal strength. Genes Dev 28, 305–316 (2014). 10.1101/gad.235473.113

37 Scatolin, G. N. et al. Single-cell transcriptional landscapes of bovine peri-implantation development. iScience 27, 109605 (2024). 10.1016/j.isci.2024.109605

38 Ramos-Ibeas, P. et al. Pluripotency and X chromosome dynamics revealed in pig pre-gastrulating embryos by single cell analysis. Nat Commun 10, 500 (2019). 10.1038/s41467-019-08387-8

39 Nowotschin, S. et al. The emergent landscape of the mouse gut endoderm at single-cell resolution. Nature 569, 361–367 (2019). 10.1038/s41586-019-1127-1

40 Cheng, S. et al. Single-Cell RNA-Seq Reveals Cellular Heterogeneity of Pluripotency Transition and X Chromosome Dynamics during Early Mouse Development. Cell Rep 26, 2593–2607.e2593 (2019). 10.1016/j.celrep.2019.02.031

41 Thowfeequ, S. et al. An integrated approach identifies the molecular underpinnings of murine anterior visceral endoderm migration. Dev Cell 59, 2347–2363.e2349 (2024). 10.1016/j.devcel.2024.05.014

42 Ross, C. & Boroviak, T. E. Origin and function of the yolk sac in primate embryogenesis. Nat Commun 11, 3760 (2020). 10.1038/s41467-020-17575-w

43 Kumar, S. et al. TimeTree 5: An Expanded Resource for Species Divergence Times. Mol Biol Evol 39 (2022). 10.1093/molbev/msac174

44 Boroviak, T. et al. Single cell transcriptome analysis of human, marmoset and mouse embryos reveals common and divergent features of preimplantation development. Development 145 (2018). 10.1242/dev.167833

45 Jin, S., Plikus, M. V. & Nie, Q. CellChat for systematic analysis of cell-cell communication from single-cell transcriptomics. Nat Protoc 20, 180–219 (2025). 10.1038/s41596-024-01045-4

46 Sozen, B., Cornwall-Scoones, J. & Zernicka-Goetz, M. The dynamics of morphogenesis in stem cell-based embryology: Novel insights for symmetry breaking. Dev Biol 474, 82–90 (2021). 10.1016/j.ydbio.2020.12.005

47 Winnier, G., Blessing, M., Labosky, P. A. & Hogan, B. L. Bone morphogenetic protein-4 is required for mesoderm formation and patterning in the mouse. Genes Dev 9, 2105–2116 (1995). 10.1101/gad.9.17.2105

48 Ying, Y. & Zhao, G. Q. Cooperation of endoderm-derived BMP2 and extraembryonic ectoderm-derived BMP4 in primordial germ cell generation in the mouse. Dev Biol 232, 484–492 (2001). 10.1006/dbio.2001.0173

49 Hadas, R. et al. Temporal BMP4 effects on mouse embryonic and extraembryonic development. Nature 634, 652–661 (2024). 10.1038/s41586-024-07937-5

50 Donnison, M. et al. Loss of the extraembryonic ectoderm in Elf5 mutants leads to defects in embryonic patterning. Development 132, 2299–2308 (2005). 10.1242/dev.01819

51 Liu, P. et al. Requirement for Wnt3 in vertebrate axis formation. Nat Genet 22, 361–365 (1999). 10.1038/11932

52 Andre, P., Song, H., Kim, W., Kispert, A. & Yang, Y. Wnt5a and Wnt11 regulate mammalian anterior-posterior axis elongation. Development 142, 1516–1527 (2015). 10.1242/dev.119065

53 Sun, X., Meyers, E. N., Lewandoski, M. & Martin, G. R. Targeted disruption of Fgf8 causes failure of cell migration in the gastrulating mouse embryo. Genes Dev 13, 1834–1846 (1999). 10.1101/gad.13.14.1834

54 Mesnard, D., Guzman-Ayala, M. & Constam, D. B. Nodal specifies embryonic visceral endoderm and sustains pluripotent cells in the epiblast before overt axial patterning. Development 133, 2497–2505 (2006). 10.1242/dev.02413

55 Schier, A. F. & Shen, M. M. Nodal signalling in vertebrate development. Nature 403, 385–389 (2000). 10.1038/35000126

56 Brumm, A. S. et al. Initiation and maintenance of the pluripotent epiblast in pre-implantation human development is independent of NODAL signaling. Dev Cell 60, 174–185.e175 (2025). 10.1016/j.devcel.2024.10.020

57 Komor, A. C., Kim, Y. B., Packer, M. S., Zuris, J. A. & Liu, D. R. Programmable editing of a target base in genomic DNA without double-stranded DNA cleavage. Nature 533, 420–424 (2016). 10.1038/nature17946

58 Wei, Q. et al. Bovine lineage specification revealed by single-cell gene expression analysis from zygote to blastocyst. Biol Reprod 97, 5–17 (2017). 10.1093/biolre/iox071

59 Blakeley, P. et al. Defining the three cell lineages of the human blastocyst by single-cell RNA-seq. Development 142, 3151–3165 (2015). 10.1242/dev.123547

60 Brennan, J. et al. Nodal signalling in the epiblast patterns the early mouse embryo. Nature 411, 965–969 (2001). 10.1038/35082103

61 Yang, R. et al. Amnion signals are essential for mesoderm formation in primates. Nat Commun 12, 5126 (2021). 10.1038/s41467-021-25186-2

62 Sasaki, K. et al. The Germ Cell Fate of Cynomolgus Monkeys Is Specified in the Nascent Amnion. Dev Cell 39, 169–185 (2016). 10.1016/j.devcel.2016.09.007

63 Xiang, L. et al. A developmental landscape of 3D-cultured human pre-gastrulation embryos. Nature 577, 537–542 (2020). 10.1038/s41586-019-1875-y

64 Pérez-Gómez, A. et al. The role of TEAD4 in trophectoderm commitment and development is not conserved in non-rodent mammals. Development 151 (2024). 10.1242/dev.202993

65 Martínez de Los Reyes, N., et al. GATA3 is not required for sheep trophectoderm development, but it plays a role in post-hatching epiblast survival. Reproduction 170 (2025). 10.1530/REP-25-0113

66 Ramos-Ibeas, P. et al. Embryonic disc formation following post-hatching bovine embryo development in vitro. Reproduction 160, 579–589 (2020). 10.1530/REP-20-0243

