## Supplementary Fig. for "A Single-Cell Transcriptomic Atlas of Symmetry Breaking Across Eutherian Mammals"

**González-Brusi et al.**

### Description of Additional Supplementary Files

**File Name:** Supplementary Data 1
**Description:** Sheep *in vivo* embryos and sequenced cells information.

**File Name:** Supplementary Data 2
**Description:** A referenced list of markers used for cell type identification.

**File Name:** Supplementary Data 3
**Description:** Top markers of cell types identified in this paper. The differential expression analysis presented was performed using the *FindAllMarkers* function from the Seurat package which uses a two-sided Wilcoxon Rank Sum test. P-values were adjusted for multiple testing using the Bonferroni correction method.

**File Name:** Supplementary Data 4
**Description:** Differentially expressed genes (DEGs) between mature and immature trophectoderm populations (Supplementary Fig. 3a). Genes were considered as DEGs if logFC > 2, adjusted p value < 0.01, expression ≥ 30% of cells within the cluster, and expression in ≤ 30% of cells in all other clusters. Gene Ontology (GO) and KEGG pathway analyses of cluster-specific markers were performed with *clusterProfiler* (v 4.16.0). Briefly, gene symbols corresponding to cluster markers were converted to Entrez IDs using the *bitr* function and analyzed with *compareCluster* function under default parameters. GO terms and KEGG pathways with an adjusted p-value < 0.01 were considered significantly enriched.

**File Name:** Supplementary Data 5
**Description:** Differentially expressed genes (DEGs) between distal parietal hypoblast, proximal parietal hypoblast, and visceral hypoblast populations (Supplementary Fig. 3a). Genes were considered as DEGs if logFC > 2, adjusted p value < 0.01, expression ≥ 30% of cells within the cluster, and expression in ≤ 30% of cells in all other clusters. Gene Ontology (GO) and KEGG pathway analyses of cluster-specific markers were performed with *clusterProfiler* (v 4.16.0). Briefly, gene symbols corresponding to cluster markers were converted to Entrez IDs using the *bitr* function and analyzed with *compareCluster* function under default parameters. GO terms and KEGG pathways with an adjusted p-value < 0.01 were considered significantly enriched.

**File Name**: Supplementary Data 6

**Description:** Differentially expressed genes (DEGs) between populations identified in gastrulating embryonic discs (Figure 2a). Genes were considered as DEGs if logFC > 2, adjusted p value < 0.01, expression ≥ 30% of cells within the cluster, and expression in ≤ 30% of cells in all other clusters. Gene Ontology (GO) and KEGG pathway analyses of cluster-specific markers were performed with *clusterProfiler* (v 4.16.0). Briefly, gene symbols corresponding to cluster markers were converted to Entrez IDs using the *bitr* function and analyzed with *compareCluster* function under default parameters. GO terms and KEGG pathways with an adjusted p-value < 0.01 were considered significantly enriched.

**File Name**: Supplementary Data 7

**Description:** Differentially expressed genes (DEGs) between anterior visceral hypoblast (AVH) 1, AVH 2, and posterior visceral hypoblast (PVH) clusters identified in E11.5 sheep embryos (Figure 3b). Genes were considered as DEGs if logFC > 2, adjusted p value < 0.01, expression ≥ 30% of cells within the cluster, and expression in ≤ 30% of cells in all other clusters. Gene Ontology (GO) and KEGG pathway analyses of cluster-specific markers were performed with *clusterProfiler* (v 4.16.0). Briefly, gene symbols corresponding to cluster markers were converted to Entrez IDs using the *bitr* function and analyzed with *compareCluster* function under default parameters. GO terms and KEGG pathways with an adjusted p-value < 0.01 were considered significantly enriched.

### Supplementary methods

#### *Inter-species comparisons*

The transcriptomes of pre-primitive streak (PS) and early-PS embryos from other mammalian species were also analysed. Either *Seurat* or *h5ad* AnnData objects were obtained from publicly available datasets for pig [1,2], marmoset [3], rabbit [4], and mouse [5-8]. Further information on methods for inter-species comparisons is provided in the Supplementary Information.

*Pig*

For the pre-PS pig embryos, the original gene matrix was annotated with an older Ensembl release that did not include some genes involved in key signalling pathways such as *NODAL*, requiring reanalysis. FASTQ files from E11 embryonic disc cells were retrieved from the EBI repository and processed with *kallisto* using a 31 k-mer size length transcriptome index built from the latest *Sus scrofa* reference genome (Ensembl release 115). Count matrices were imported with *tximport* [9] and transformed into a *Seurat* object. Independent processing identified 42 epiblast (EPI) and 18 visceral hypoblast (VH) cells, consistent with the original annotation. No asymmetry was detected in either lineage, as assessed by *TBXT* expression in the EPI and *CER1, LEFTY1, LEFTY2, HHEX, CER1, GSC,* and *FZD5* expression in the VH.

For early PS pig embryos [2], only E11.5 embryos were retained. The original annotation was adjusted as follows: "Trophoblast" -> "TE", "ExE Endoderm" -> "PH", "Gut/Hypoblast" -> "PVH", "Anterior Hypoblast/AVE" -> "AVH", "Epiblast 1", "Epiblast 3" and "Epiblast 4" -> "EPI", "Anterior Primitive Streak/Node" -> "APS", "Primitive Streak 2", "Primitive Streak 3" -> "PS", "Nascent Mesoderm 1", "Nascent Mesoderm 2" -> "nMeso". Cells in AVH and PVH clusters were retained if they expressed at least one VH marker (*NODAL, OTX2, LHX1, HHEX*).

*Cow*

For the cow dataset [10], D12 and D14 sample expression matrices and cell annotations were downloaded from the GEO repository (GSE234335). Due to annotation errors affecting key genes (*LEFTY1, LEFTY2*), reannotation was performed using the original Ensembl v109 biomaRt dataset. Briefly, genes encoding high-confidence human homologs whose gene names differed from those in the original feature matrix were reannotated, while the remaining annotations were retained unchanged. Cells with fewer than 80,000 counts, between 2,000 and 6,000 detected features and less than 20% mitochondrial counts were retained for downstream analyses. Each sample was processed independently using the following parameters: *FindVariableFeatures* with 2000 features, *ScaleData* regressing with nFeatureRNA, mitochondrial count percentage and cell cycle S and G2M phases scores. *RunPCA* was performed using 100 PCs, *RunUMAP* using the first 50 dimensions, *FindNeighbors* using the first 30 dimensions and *FindClusters* with a resolution of 0.2.

D12 samples were integrated to generate the pre-PS subset. Integration was performed using reciprocal PCA via *IntegrateLayers* following individual sample merging. *RunUMAP* and *FindNeighbors* were run using the first 30 dimensions, and *FindClusters* with a resolution of 0.2. Lineage segregation was assessed by visualizing established key markers, consistent with those used in our dataset, using *FeaturePlot*. Original clusters 0 and 3 (distal parietal hypoblast) were excluded. Cluster 2 was further subset and subclustered at a resolution of 0.3 to distinguish PH, PVH and AVH. The resulting annotations were transferred back to the main Seurat object.

D14 samples following the same strategy as for D12 samples to generate the early-PS subset. Annotation was performed using the same key markers defined in sheep and visualized using *FeaturePlot*. Unsupervised clustering identified TE, dPH (removed), PH, VH and EPI-PS-nMeso. VH cells were subset and subclustered at the default resolution to discriminate PVH and AVH. Separation of EPI-PS-nMeso populations required increasing the clustering resolution to 1.2.

*Marmoset*

For the marmoset dataset [3], Carnegie stage (CS) 5 and CS6 embryos were retained for pre-PS and early-PS stages, respectively, using the original annotation. Maternal cells, primordial germ cells (PGCs), and stalk were excluded. In CS5 embryos, the amnion did not participate in inter-lineage communication, probably due to the reduced cell number and was thus excluded from the analysis. Within CS6, the ED cluster was subclustered using *FindClusters* (resolution = 1.7). The SOX2+ subcluster was annotated as “EPI”, while the reminder of the ED cluster was annotated as “PS”. Visceral endoderm (VE) cells were subsetted and subclustered (*FindClusters*, resolution = 0.8), yielding anterior visceral hypoblast (AVH, validated by *CER1, FZD5, GSC* or *HHEX* expression) and posterior visceral hypoblast (PVH) populations.

*Rabbit*

For the rabbit dataset [4], stages 0 and 3 were selected. Stage 0 embryos, lacking PS, were developmentally comparable to sheep pre-PA E11 embryos. Stage 1 and 2 embryos contained only a small number of PS cells (*TBXT+*, *MIXL1+*) and no nascent mesoderm cells (*TBX6+, BMP4+*), and were excluded. Stage 3 embryos resembled sheep early-PS E11.5 embryos, with additional *TFAP2C+* cells within the ED, identified as emerging PGCs, and two extraembryonic hypoblast subpopulations (parietal hypoblast, PH; yolk sac endoderm, YSE). *TFAP2C+* ED cells, PGCs, and

PH were removed, but YSE was retained, as its transcriptome closely resembled proximal parietal hypoblast in pig and sheep.

Stage 0 and 3 datasets were scaled using cell-cycle scores, embryo identity, amplification batch, sequencing batch, and expression of forbidden genes list from the original study as regression variables. Dimensionality reduction was performed with *RunPCA* (100 PCs) and *RunUMAP* (25 dimensions; min.dist = 0.2; n.neighbors = 30). Clustering was performed with *FindNeighbors* (25 dimensions) and *FindClusters* (resolution = 0.5). Embryo "Rab1_f1e4" was excluded due to failed integration. Original gene name annotations were standardized by removing mm9 and hg19 orthology labels to facilitate cross-species comparisons.

For stage 3 dataset, clusters corresponding to YSE-VE and PS-APS-nMeso (idents 5 and 4) were further subclustered using *FindClusters* (resolutions = 1 and 0.5, respectively). Subclusters were annotated based on markers expression: *AFP+/RSPO3+* YSE, *AFP-/RSPO3+* PVH, *FZD5+/CER1+* AVH (hypoblast lineages), *TBX6+* nMeso, *FOXA2+* APS, *TBXT+TBX6-FOXA2-* PS (PS-derived lineages).

*Mouse*

Mouse datasets were processed as follows for integration into two stages (pre-PS: E5.25-5.5; early-PS: E6.25-E6.5):

- GSE109071 [7]: FASTQ files from E5.25, E5.5, E6.25 and E6.5 cells obtained with the smartseq2 protocol were retrieved from the EBI repository and processed with *kallisto* using a 17 k-mer size length transcriptome index built from the latest *Mus musculus* reference genome (Ensembl release 115). Count matrices were imported with *tximport* [72] and transformed into a *Seurat* object. Both pre-PS and early-PS embryos were scaled using cell-cycle scoring, mitochondrial transcript counts, and embryo identity.
- GSE123046 [5]: FASTQ files from E5.5 and E6.5 cells obtained with the 10x Chromium v2 protocol were retrieved from the EBI repository and processed with the kb-python wrapper of *kallisto bustools* [11] using a 17 k-mer size length transcriptome index built from the latest *Mus musculus* reference genome (Ensembl release 115). A Seurat object corresponding to each individual sample was created from its count matrix with the *CreateSeuratObject* function. Cells with a RNA count lower than 5600, less than 1000 detected genes or more than 5% mitochondrial mRNA percent were filtered out. Both pre-PS and early-PS embryos were scaled using cell-cycle scoring and mitochondrial transcript counts percentageDatasets from each time point were integrated using RPCA.
- Thowfeequ et al. 2024 [8]: AnnData objects were downloaded from the ScialdoneLab GitHub repository and converted to *Seurat* objects.

The final pre-PS *Seurat* object for cell signaling analysis was a product of integrating E5.25-5.5 cells from the datasets mentioned above using CCA. *RunPCA* was executed on 100 dimensions, *FindNeighbors* on 50 dimensions, *FindClusters* at a resolution of 0.2, and *RunUMAP* on 50 dimensions with *min.dist* set to 0.3. A *Bmp4*- ExE population corresponding to the ectoplacental cone (EPC) was excluded from further analysis.

Primitive endoderm populations were subclustered with 0.4 resolution and labeled according to gene markers: *Epas1+* for Parietal endoderm (PE), *Apln+* for exVE, *Eomes+/Nodal+* for emVE (both anterior and posterior), and *Lefty1+/Cer1+* for AVE/DVE (termed AVE for simplification).

**Supplementary Table 1.** Details of primers used for single guide RNA (sgRNA) production, and for embryo genotyping.

| ID | Sequence (5´→3´) | Use |
|---|---|---|
| T7 Guide-it ovNODAL | CCTCTAATACGACTCACTATAGGAGCCCACCAGGCGTGCAAGGTTTAAGAGCTATGC | To produce gRNA against ovine *NODAL*. Target sequence is underlined. |
| Geno ovNODAL Fw | GGGTGGGGTAGGGGTGTTAT | To genotype ovine embryos by Sanger. |
| Geno ovNODAL Rv | TCCTCAAATGGCAAGTCCCC | |

**Supplementary Table 2.** Embryo transfers performed and CRISPR-Cas9 efficiency. Day (D) 7 embryos from the *NODAL*-targeted (BE+G; microinjected with *NODAL* gRNA and BE3 mRNA) group and control (BE; microinjected with BE3 mRNA only) were transferred to three ewes. Embryos were recovered at embryonic day (E) 12 (after 5 days) and 14 (after 7 days). Embryo recovery efficiency, genotyping efficiency, editing efficiency, as well as knock-out (KO), edited non-KO (ENK) and wild-type (WT) rates are shown.

| Embryonic day (E) | Embryos transferred | Embryos recovered (%) | Genotyped embryos (%) | Edited embryos (%) | KO embryos (%) | ENK embryos (%) | WT embryos (%) |
|---|---|---|---|---|---|---|---|
| 12_ewe 1 | 46 BE+G*<br>12 BE | 36/58 (62.07) | 35/36 (97.22) | 23/35 (65.71) | 14/35 (40) | 9/35 (25.71) | 12/35 (37.5) |
| 12_ewe 2 | 18 BE+G<br>5 BE | 16/23 (69.57) | 14/16 (87.5) | 10/14 (71.43) | 4/14 (28.57) | 6/14 (42.86) | 4/14 (30.78) |
| 14 | 28 BE+G<br>12 BE | 28/40 (70.00) | 27/28 (96.43) | 18/27 (66.67) | 16/27 (59.26) | 2/27 (7.41) | 9/27 (33.33) |

*Of the 46 transferred embryos, 10 were blastocysts and 36 were morulae. The remaining transferred embryos were at the blastocyst stage.

**Supplementary Table 3**. Details of reagents used within this study (antibodies, IF, and cell culture reagents).

| Antibodies | Antigen | Host species | Dilution | Company | Cat No. | RRID |
|---|---|---|---|---|---|---|
| Primary | SOX2 | Rat | 1:100 | Invitrogen | 14-9811-80 | AB_11219471 |
| | SOX17 | Goat | 1:100 | R&D | AF1924 | AB_355060 |
| | FOXA2 | Rabbit | 1:100 | Cell Signalling | 8186S | AB_10891055 |
| | GATA3 | Rabbit | 1:100 | Abcam | ab199428 | AB_2819013 |
| | CDX2 | Mouse | 1:100 | Biogenex | MU392A-UC | AB_2923402 |
| | OTX2 | Goat | 1:100 | R&D | AF1979 | AB_2157172 |
| | BRACHYURY (T) | Goat | 1:100 | R&D | AF2085 | AB_2200235 |
| Secondary | Anti-Rat IgG 488 | Donkey | 1:300 | Life Technologies | A-21208 | AB_2535795 |
| | Anti-Goat IgG 647 | Donkey | 1:300 | Life Technologies | A-32849 | AB_2762840 |
| | Anti-Rabbit IgG 488 | Donkey | 1:300 | Life Technologies | A-32790 | AB_2762833 |
| | Anti-Rabbit IgG 555 | Donkey | 1:300 | Life Technologies | A-32794 | AB_2762833 |
| | Anti-Rabbit IgG 647 | Donkey | 1:300 | Life Technologies | A-32795 | AB_2866496 |
| | Anti-Mouse IgG 647 | Donkey | 1:300 | Life Technologies | A-32787 | AB_2762823 |
| | Phalloidin 488 | - | 1:400 | Invitrogen | A-12379 | - |
| **IF reagents** | Paraformaldehyde | | | Electron Microscopy Sciences | 15710 | |
| | Triton-X | | | Sigma-Aldrich | T9284 | |
| | Tween 20 | | | Sigma-Aldrich | P2287 | |
| | Bovine serum Albumin | | | Sigma-Aldrich | 9647 | |
| | Fluoroshield with DAPI | | | Sigma-Aldrich | F6057 | |
| **Single cell dissociation** | TrypLE express | | | Gibco | 12604-013 | |
| ***In vitro* embryo production and culture** | TCM-199 medium | | | Sigma-Aldrich | M4530 | |
| | Epidermal Growth Factor | | | Sigma-Aldrich | E4127 | |
| | Fetal Bovine Serum | | | Sigma-Aldrich | F2442 | |
| | Gentamicine sulphate | | | Sigma-Aldrich | G1272 | |
| | IVF medium | | | Stroebech media | 5.06.020 | |
| | Heparin | | | Merck | 375095 | |

| | | |
|---|---|---|
| Bovi-Pure® | Nidacon | BP-100 |
| IVC medium | Stroebech media | 5.07.020 |
| Heavy oil | Stroebech media | 6.09.050 |
| Euroflush | IMV Technologies | 019450 |
| DMEM/F12 medium | Gibco | 11320-033 |
| Neurobasal medium | Gibco | 21103-049 |
| N-2 supplement | Gibco | 17502-048 |
| B-27 supplement | Gibco | 17504-044 |
| L-Glutamine | Gibco | 25030-032 |
| Penicillin/Streptomycin | Gibco | 15140-148 |

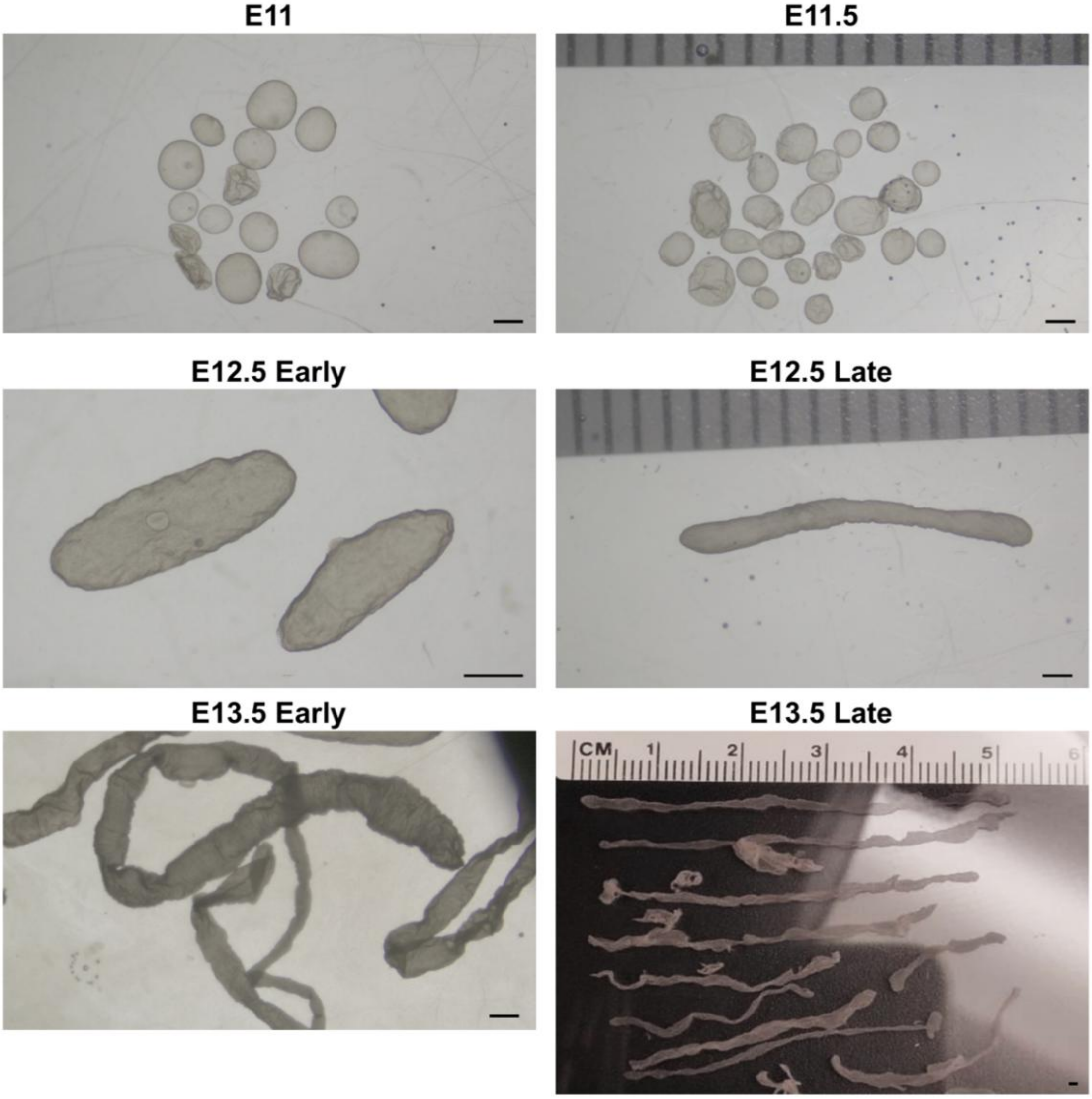


**Supplementary Figure 1. Representative images of sampled embryos**. Related to Figure 1a. See Supplementary Table 1 for sample collection details. Scale bars: 1 mm.

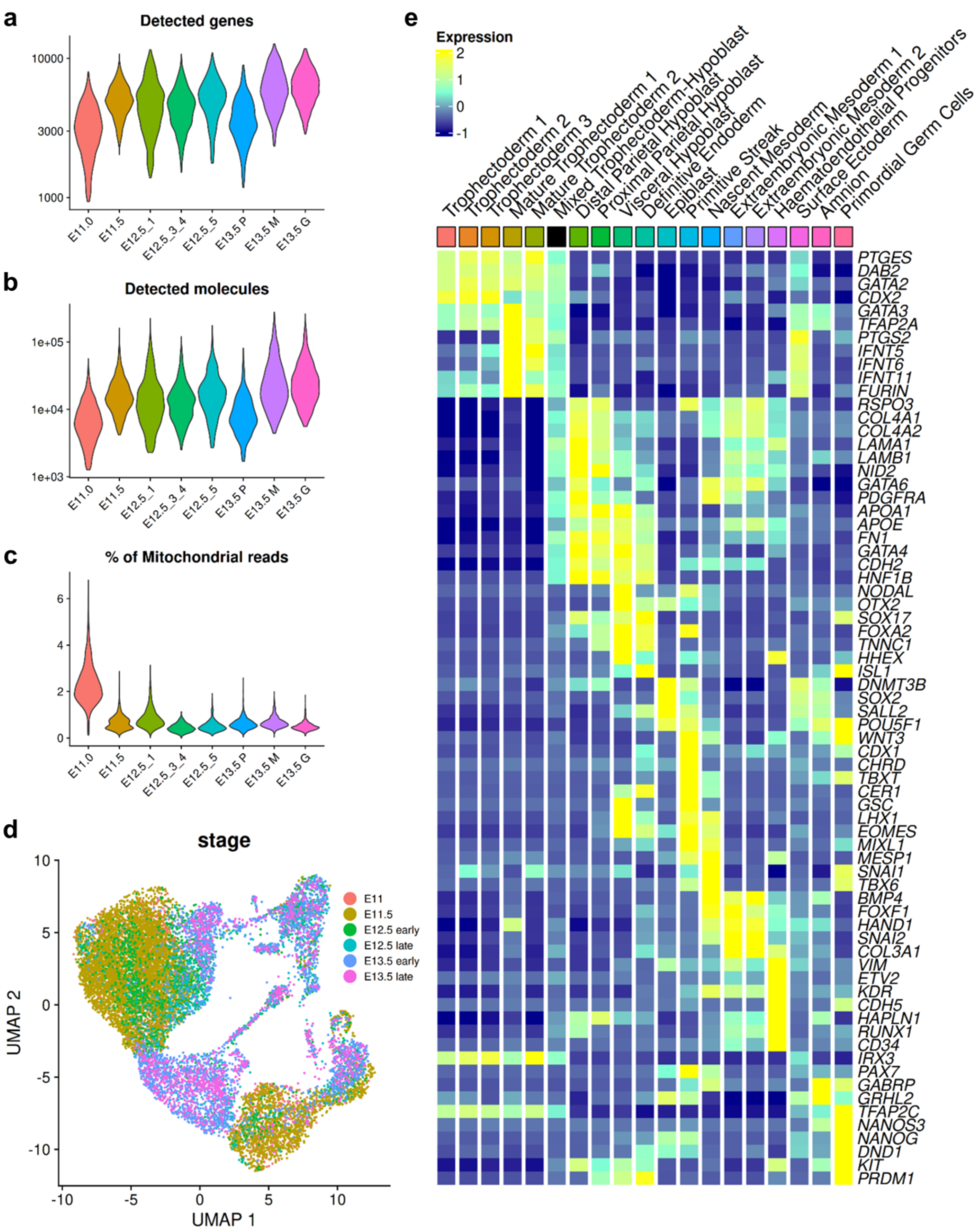


**Supplementary Figure 2. Cell distribution per sample and gene markers used to define cell identity.**

a-c) Violin plots illustrating the number of detected genes, the number of detected molecules, and the percentage of mitochondrial reads detected per cell per sample.

d) UMAP plot showing atlas cells from Figure 1b) coloured by sample.

e) Heatmap showing some of the markers used for cell type identification. A full list of these markers and the source publications are available in Supplementary Table 2.

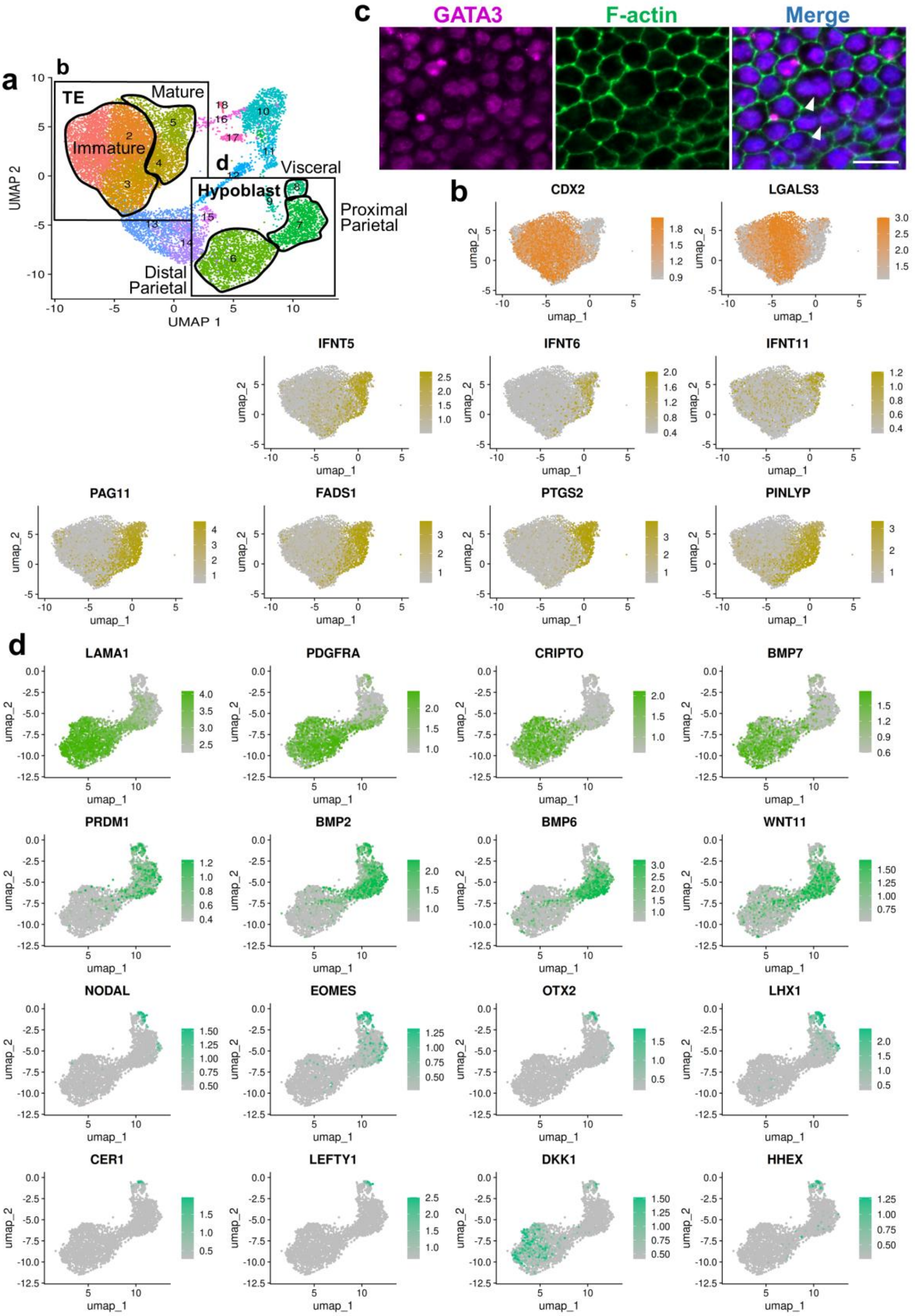


**Supplementary Figure 3. Development of extraembryonic lineages (trophectoderm and hypoblast)**.

a) UMAP plot from Figure 1b) showing trophectoderm and hypoblast populations.

b) Feature plots of the selected region in a), coloured by normalised gene expression of selected genes upregulated in more immature or mature trophectoderm cells.

c) Representative images of binucleate trophectoderm cells pointed by arrowheads in E12.5 sheep embryos. GATA3 (trophectoderm), F-actin (cellular membranes), and DAPI (nuclei) staining. Scale bar: 20 µm.

d) Feature plots of the selected region in a), coloured by normalised gene expression of selected genes upregulated in distal parietal, proximal parietal, or visceral hypoblast cells.

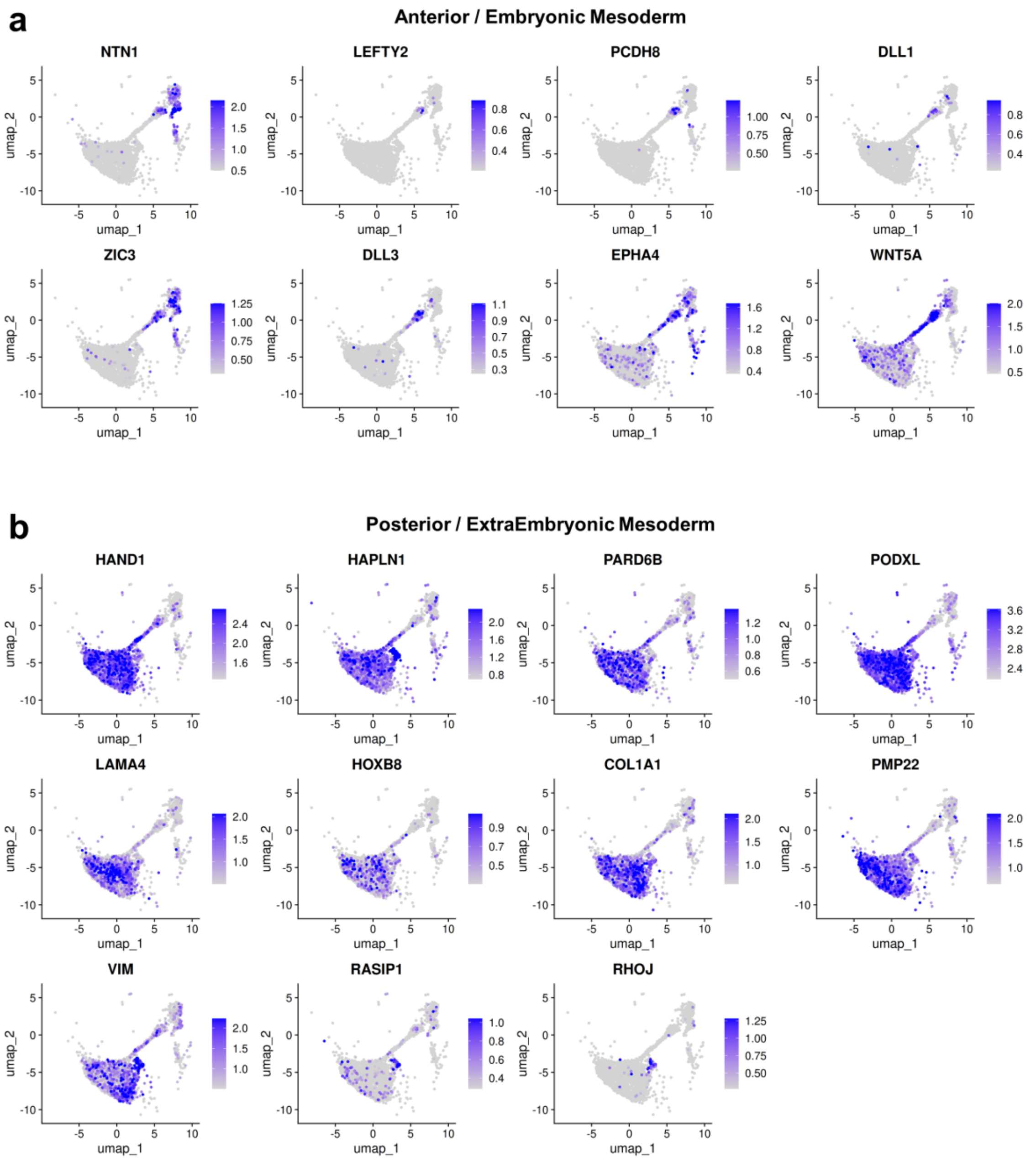


**Supplementary Figure 4. UMAP feature plots of selected a) anterior / embryonic and b) posterior / extraembryonic mesoderm marker genes.** Feature plots coloured by normalised gene expression. Related to Figure 2a. Feature plot scales were determined by the 5$^{th}$ and 95$^{th}$ percentiles.

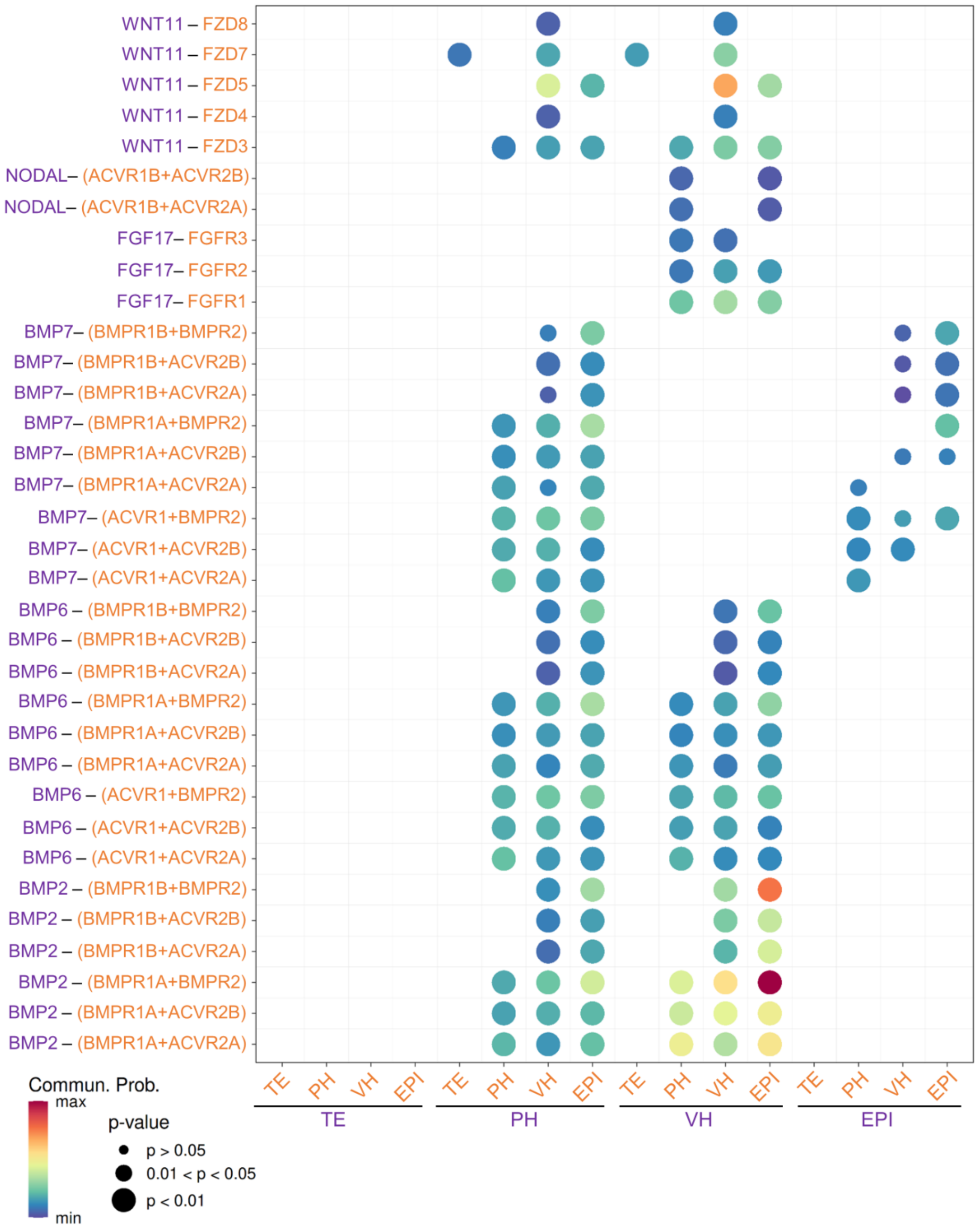


**Supplementary Figure 5. CellChat bubble plot showing significant ligand-receptor pairs that contribute to the signalling pathways mediating communication between selected lineages in pre-primitive streak (pre-PS) E11 sheep embryos.** Related to Figure 6a. Each ligand-receptor pair is colour matched to the expressing lineage on the x-axis.

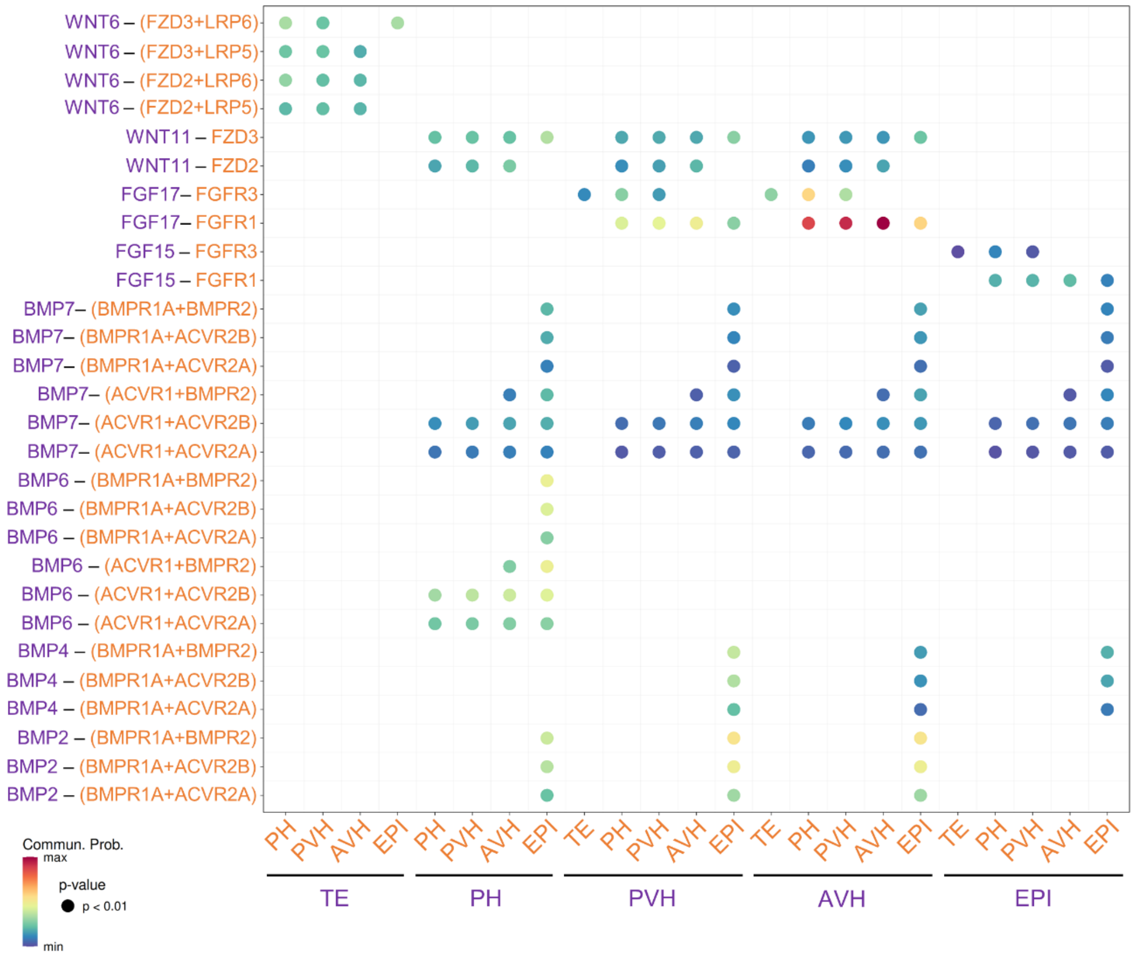


**Supplementary Figure 6. CellChat bubble plot showing significant ligand-receptor pairs that contribute to the signalling pathways mediating communication between selected lineages in pre-primitive streak (pre-PS) E12 cow embryos [10].** Related to Figure 6b. Each ligand-receptor pair is colour matched to the expressing lineage on the x-axis.

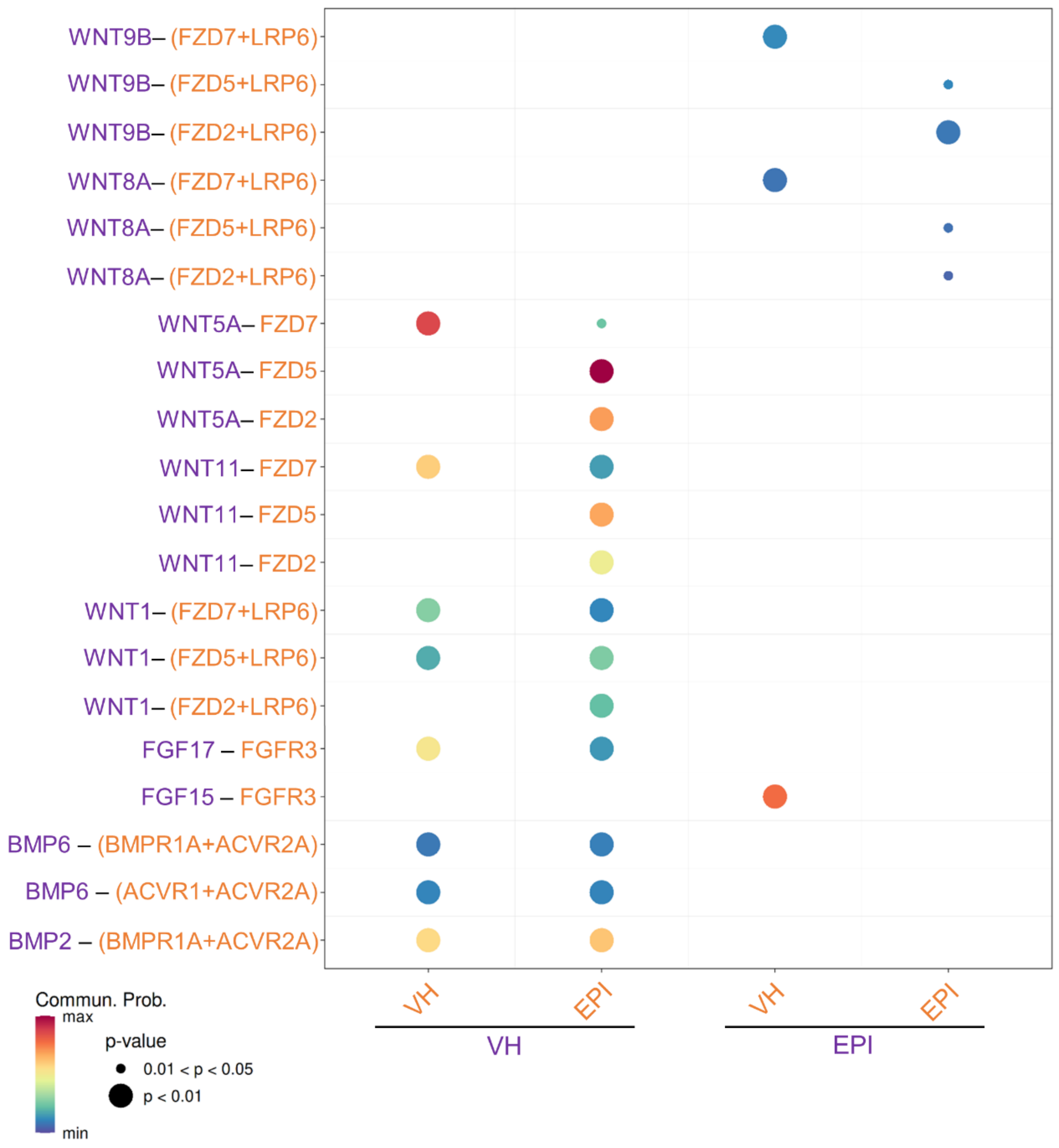


**Supplementary Figure 7. CellChat bubble plot showing significant ligand-receptor pairs that contribute to the signalling pathways mediating communication between selected lineages in pre-PS E11 pig embryos [1].** Related to Figure 6c. Each ligand-receptor pair is colour matched to the expressing lineage on the x-axis.

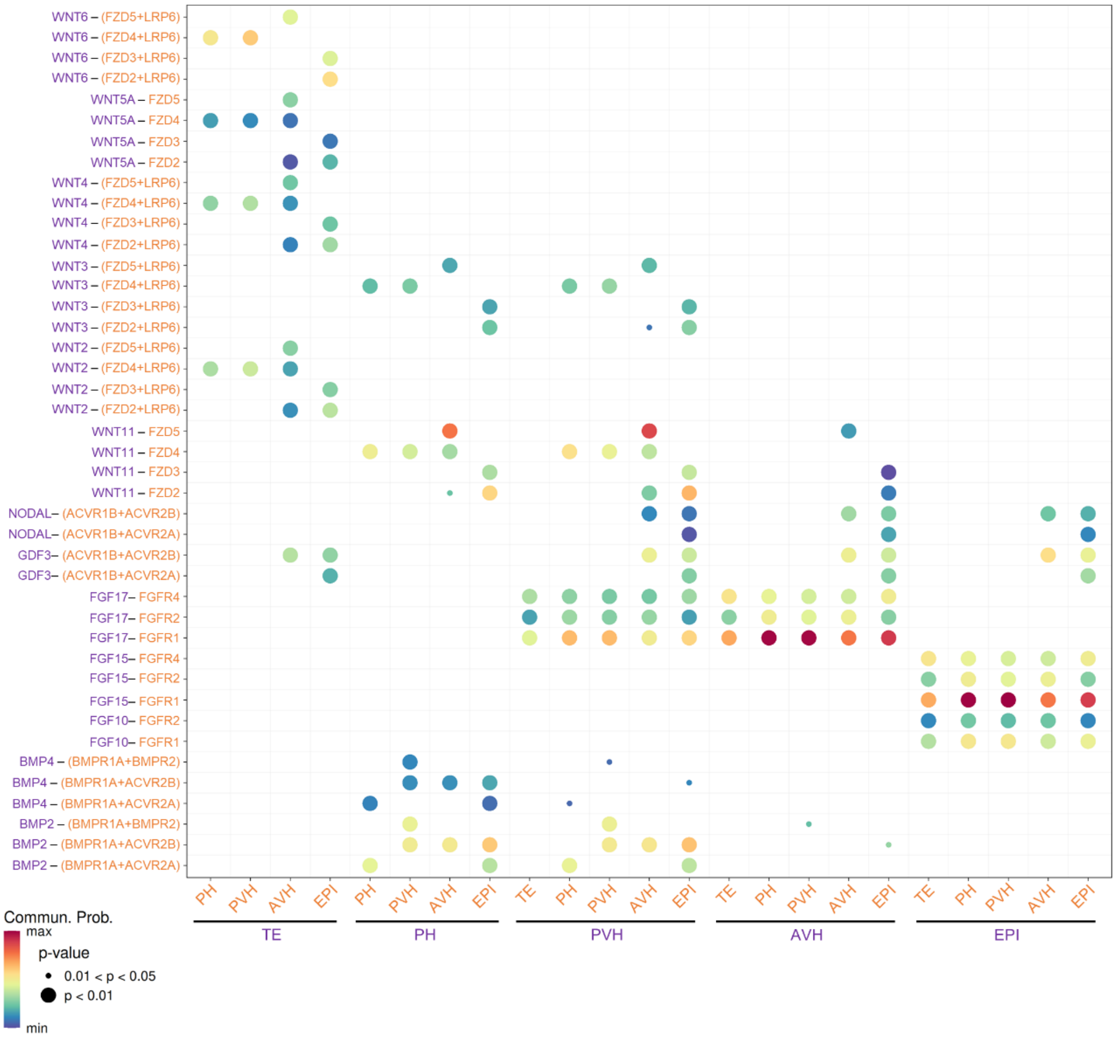


**Supplementary Figure 8. CellChat bubble plot showing significant ligand-receptor pairs that contribute to the signalling pathways mediating communication between selected lineages in pre-PS Stage 0 (E6) rabbit embryos [4].** Related to Figure 6d. Each ligand-receptor pair is colour matched to the expressing lineage on the x-axis.

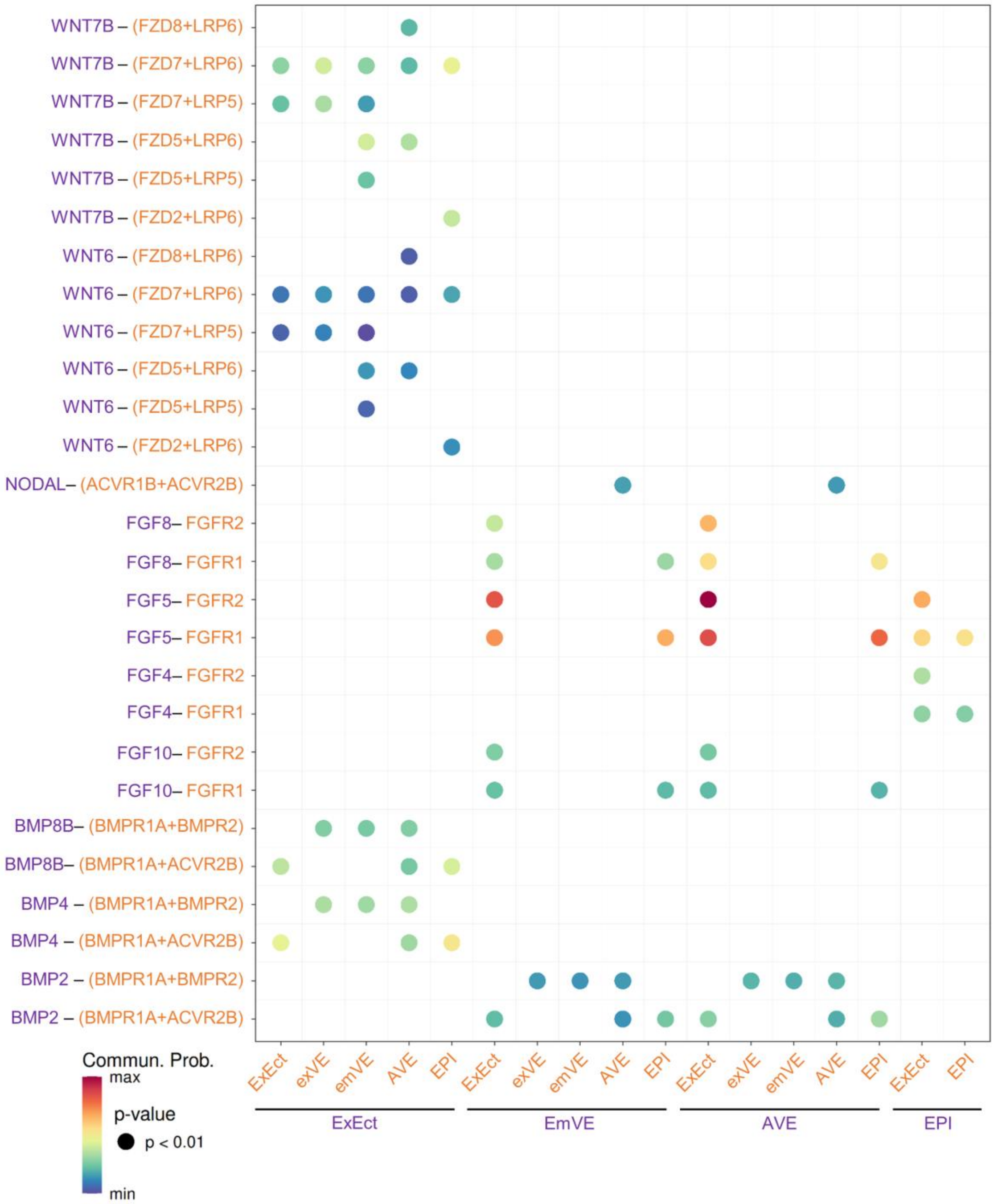


**Supplementary Figure 9. CellChat bubble plot showing significant ligand-receptor pairs that contribute to the signalling pathways mediating communication between selected lineages in pre-PS E5.25-5.5 mouse embryos [5,7,8].** Related to Figure 6e. Each ligand-receptor pair is colour matched to the expressing lineage on the x-axis.

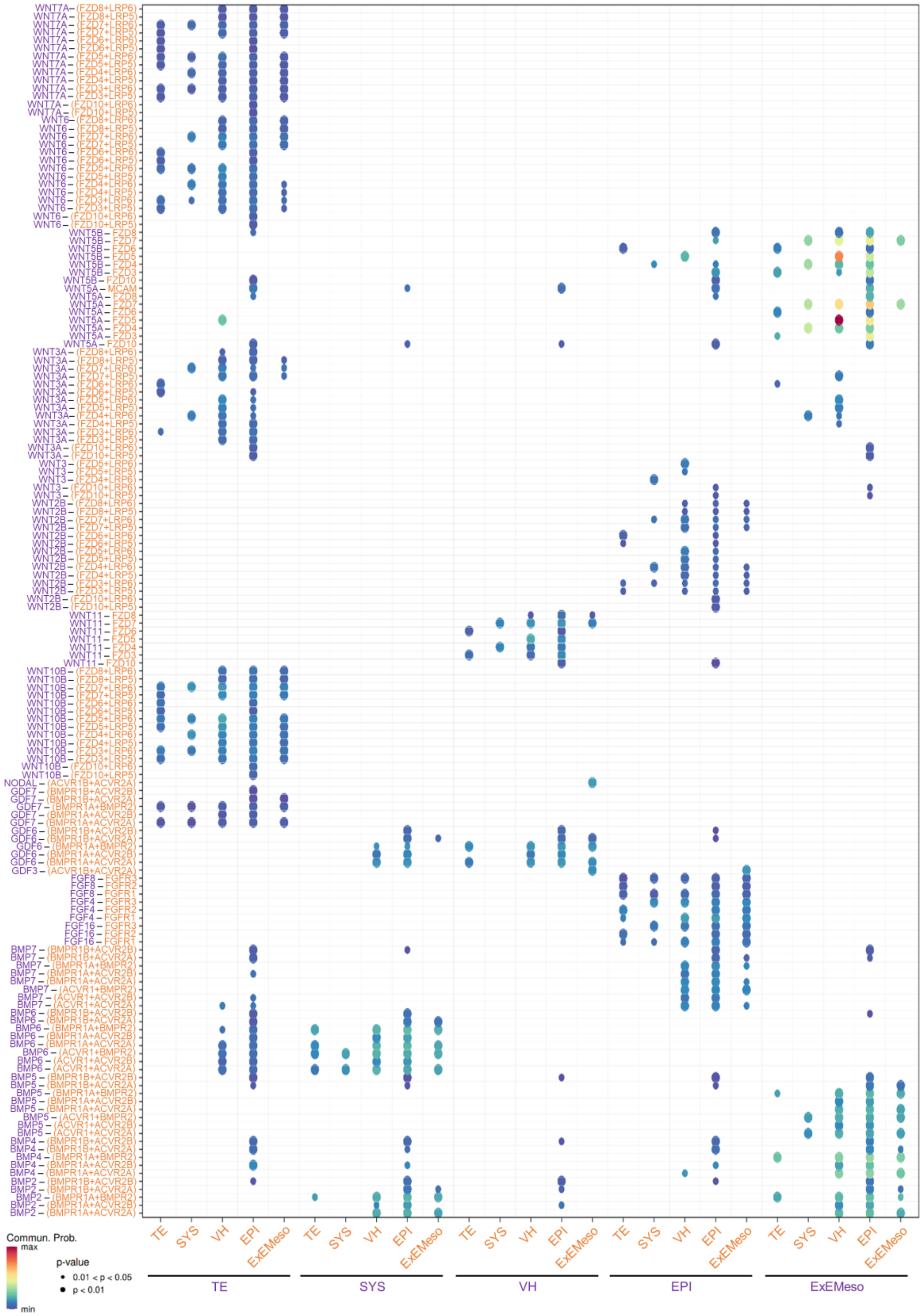


**Supplementary Figure 10. CellChat bubble plot showing significant ligand-receptor pairs that contribute to the signalling pathways mediating communication between selected lineages in**

**pre-PS CS5 (E13-15) marmoset embryos [3].** Related to Figure 6f. Each ligand-receptor pair is colour matched to the expressing lineage on the x-axis.

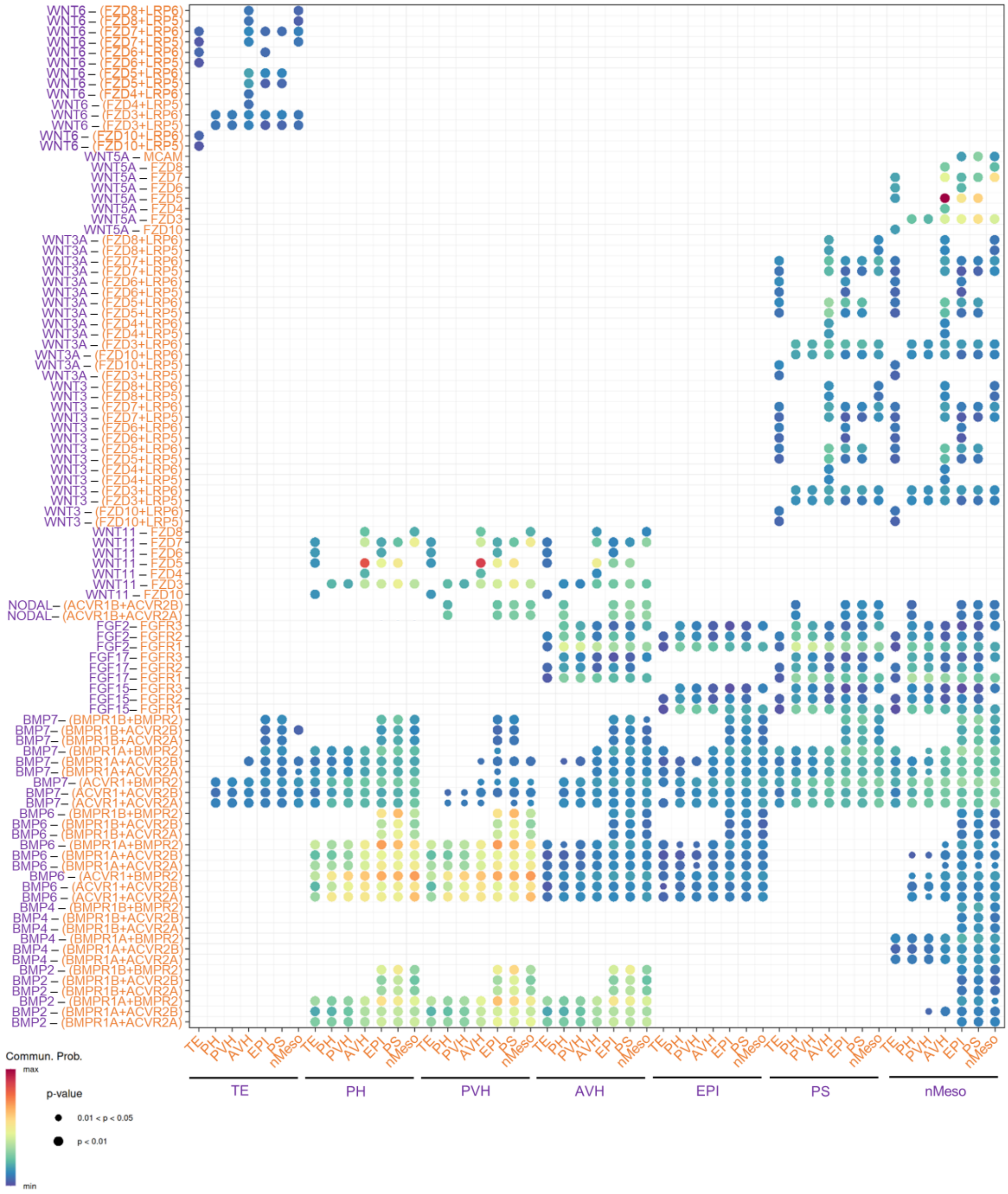


**Supplementary Figure 11. CellChat bubble plot showing significant ligand-receptor pairs that contribute to the signalling pathways mediating communication between selected lineages in early-primitive streak (early-PS) E11.5 sheep embryos.** Related to Figure 6g. Each ligand-receptor pair is colour matched to the expressing lineage on the x-axis.

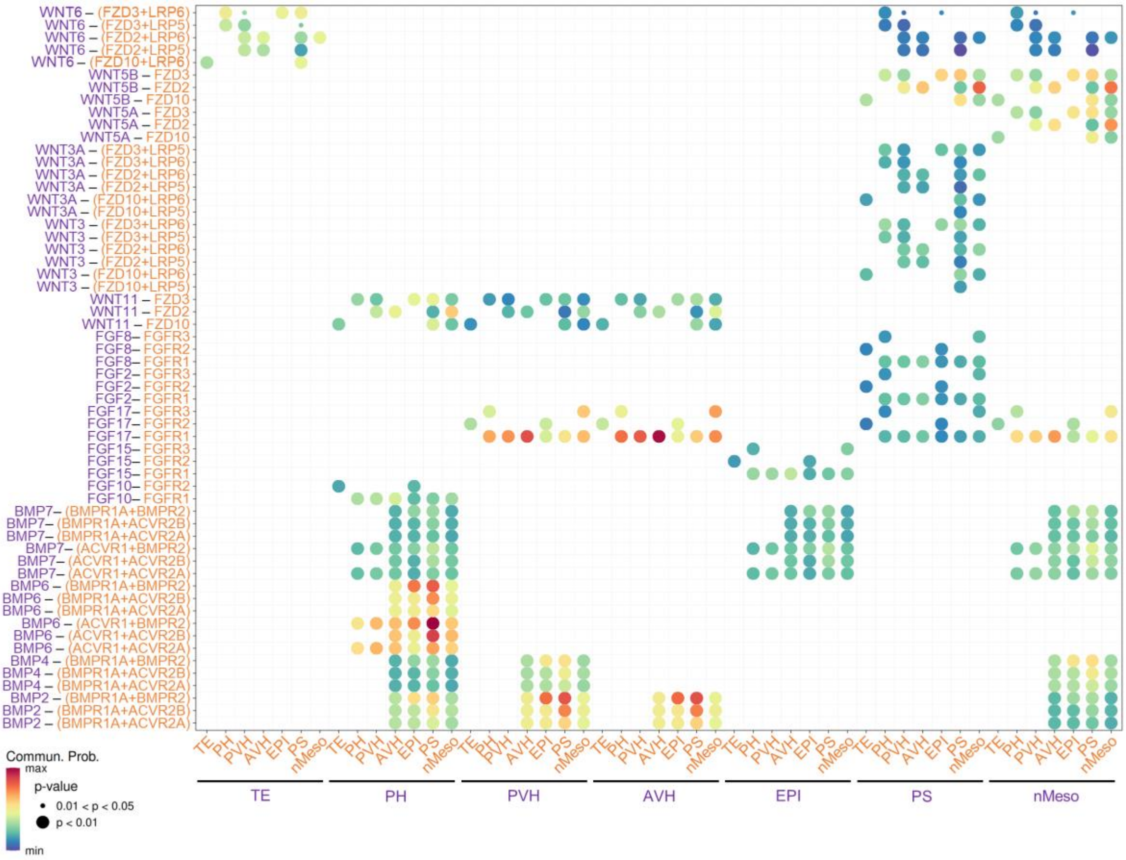


**Supplementary Figure 12. CellChat bubble plot showing significant ligand-receptor pairs that contribute to the signalling pathways mediating communication between selected lineages in early-primitive streak (early-PS) E14 cow embryos.** Related to Figure 6h. Each ligand-receptor pair is colour matched to the expressing lineage on the x-axis.

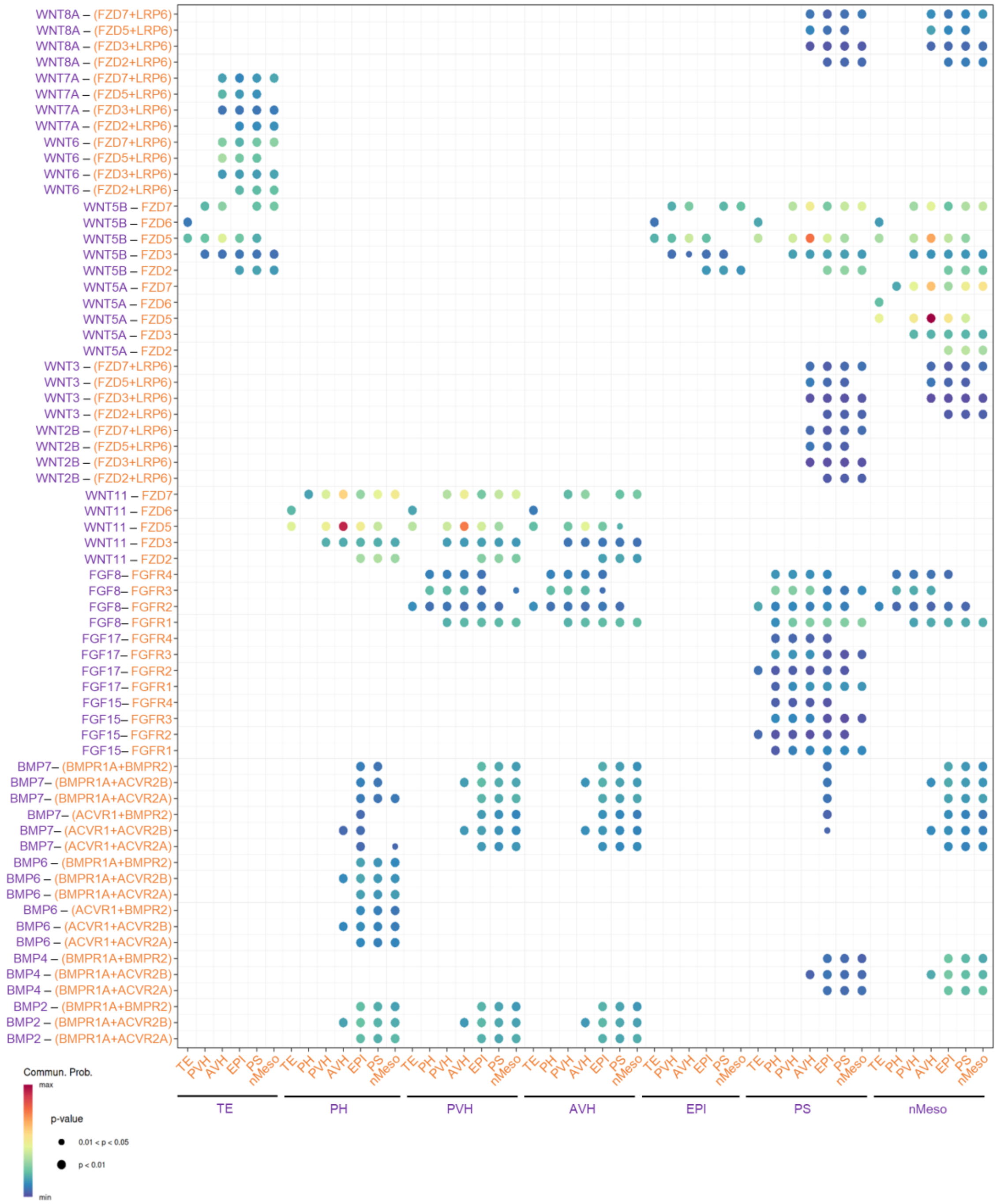


**Supplementary Figure 13. CellChat bubble plot showing significant ligand-receptor pairs that contribute to the signalling pathways mediating communication between selected lineages in early-PS E11.5 pig embryos [2].** Related to Figure 6i. Each ligand-receptor pair is colour matched to the expressing lineage on the x-axis.

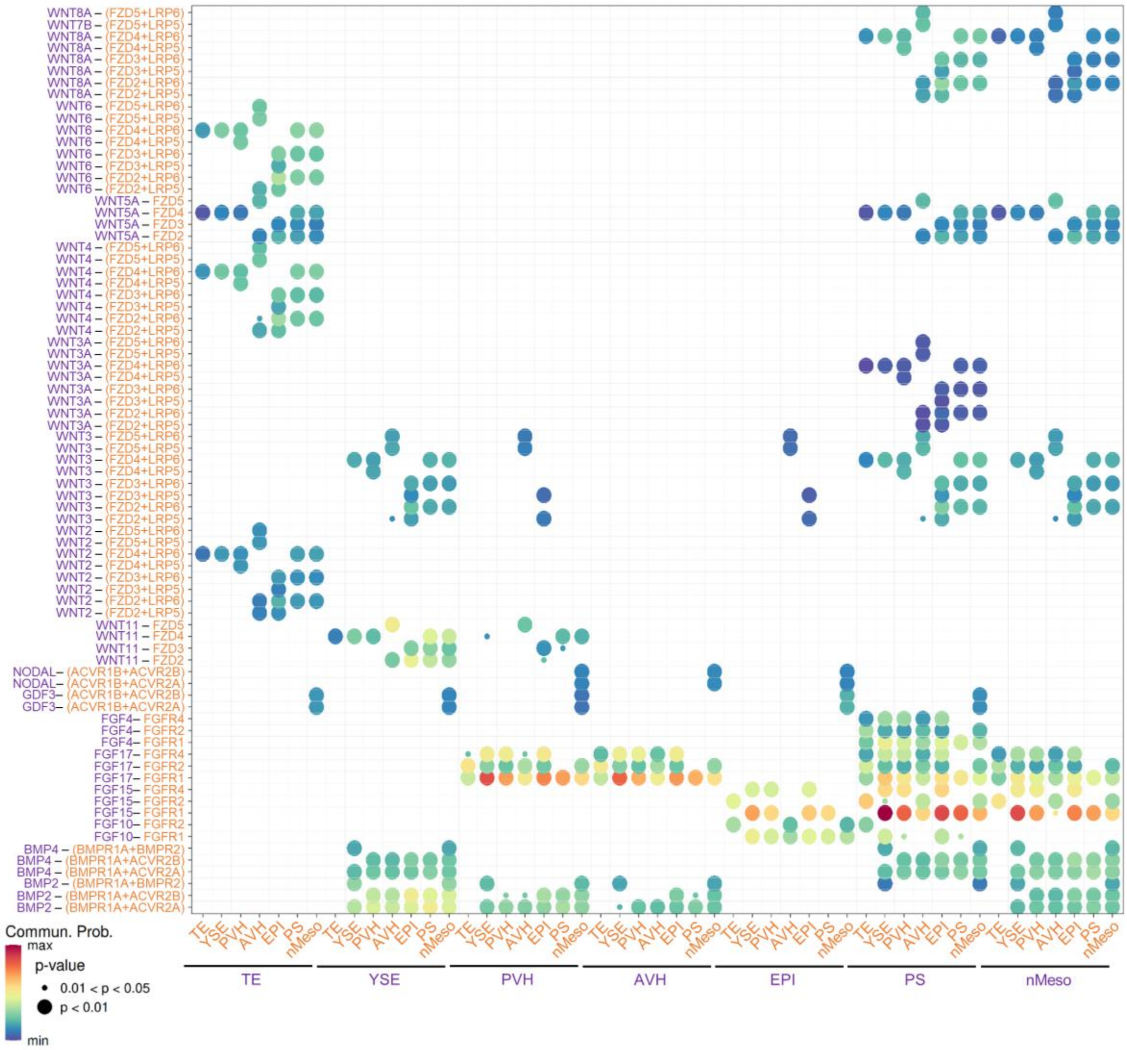


**Supplementary Figure 14. CellChat bubble plot showing significant ligand-receptor pairs that contribute to the signalling pathways mediating communication between selected lineages in early-PS Stage 3 (E6.6 and 7) rabbit embryos [4].** Related to Figure 6j. Each ligand-receptor pair is colour matched to the expressing lineage on the x-axis.

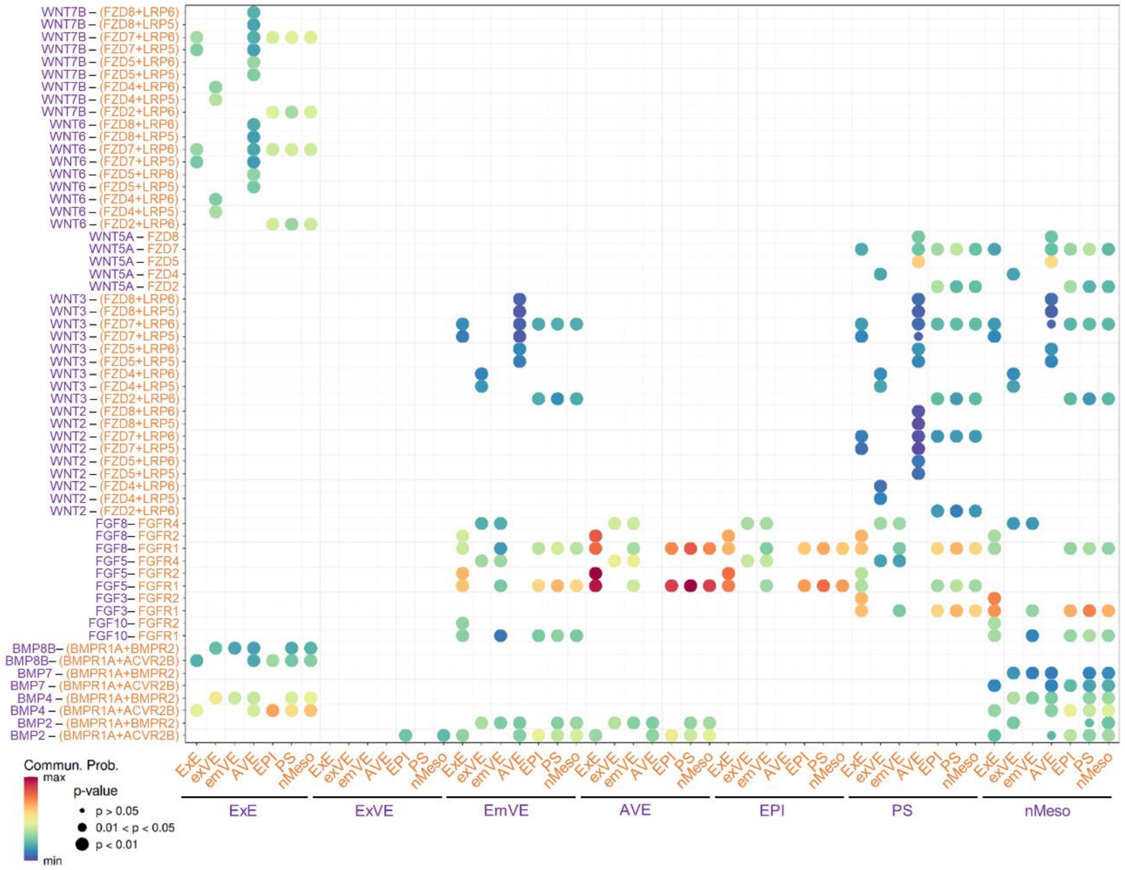


**Supplementary Figure 15. CellChat bubble plot showing significant ligand-receptor pairs that contribute to the signalling pathways mediating communication between selected lineages in early-PS E6.25-6.5 mouse embryos [5].** Related to Figure 6k. Each ligand-receptor pair is colour matched to the expressing lineage on the x-axis.

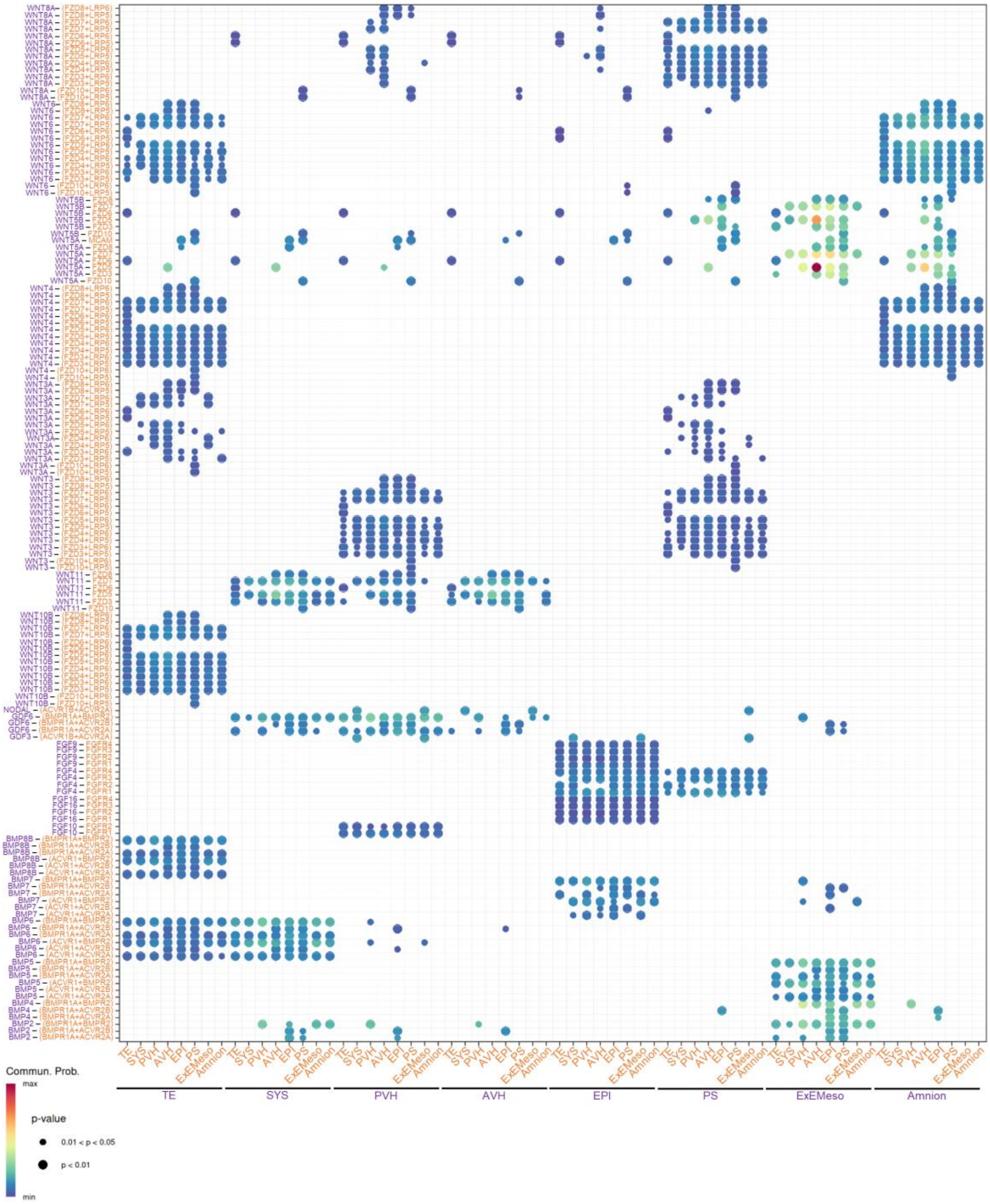


**Supplementary Figure 16. CellChat bubble plot showing significant ligand-receptor pairs that contribute to the signalling pathways mediating communication between selected lineages in early-PS CS6 (E15-17) marmoset embryos [3].** Related to Figure 6l. Each ligand-receptor pair is colour matched to the expressing lineage on the x-axis.

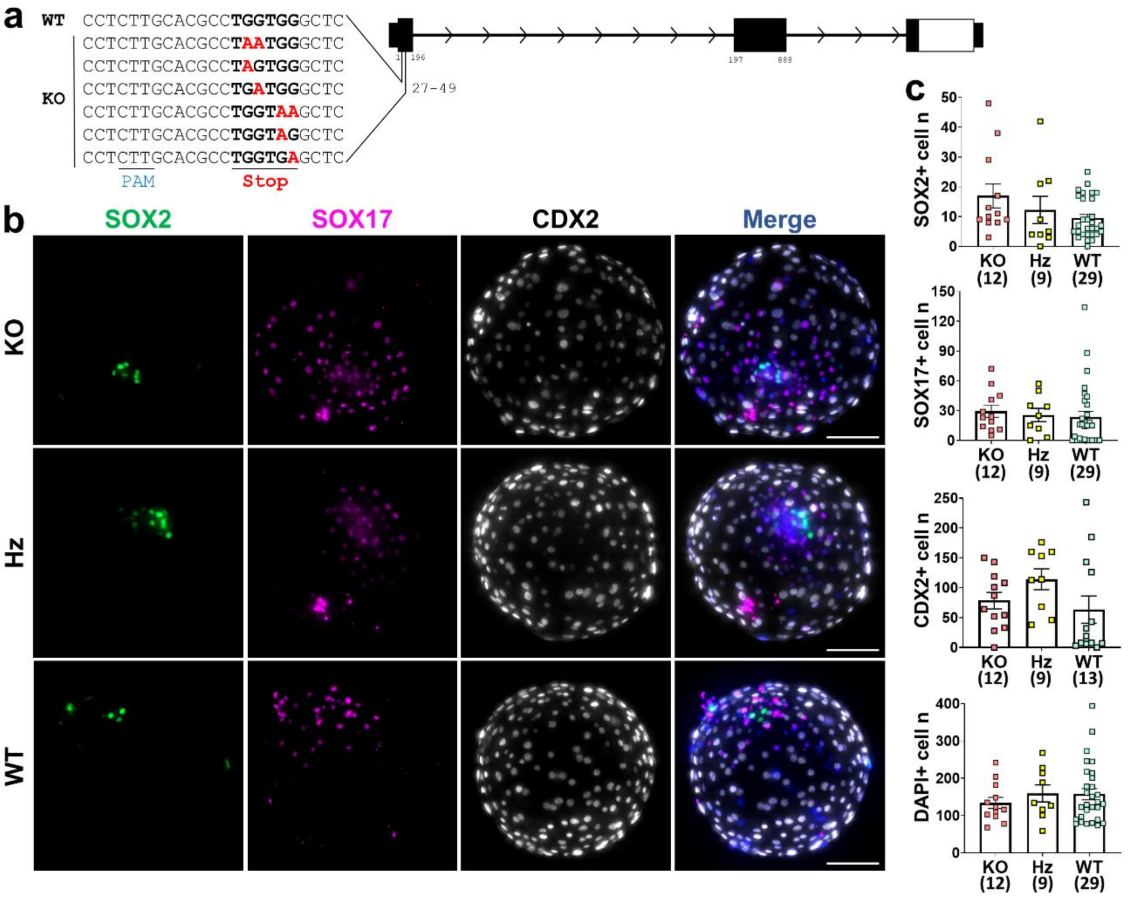


**Supplementary Figure 17. *NODAL* ablation does not impair *in vitro* sheep embryo development up to day (D) 8.**

a) Schematic representation of the *NODAL* gene. The gRNA was designed to target two consecutive TGG codons within exon 1. Red letters indicate some of the potential base substitutions leading to the generation of premature stop codons (TAA, TAG, or TGA). KO: knock-out; PAM: protospacer adjacent motif; WT: wild-type.

b) Representative immunofluorescence images of *NODAL* knock-out (KO), heterozygous (Hz) and wild-type (WT) D8 blastocysts stained for SOX2 (green; epiblast), SOX17 (magenta; hypoblast) and FOXA2 (white; trophectoderm). Nuclei were counterstained with DAPI (merge). Scale bars: 100 µm.

c) Scatter plots showing the number of SOX2+ epiblast cells, SOX17+ hypoblast cells, CDX2+ trophectoderm cells, and DAPI+ total cells in KO, Hz, and WT D8 blastocysts (mean ± s.e.m.). The number of analysed embryos is shown below columns. No statistically significant differences were detected ($P > 0.05$; non-parametric Kruskal-Wallis test).

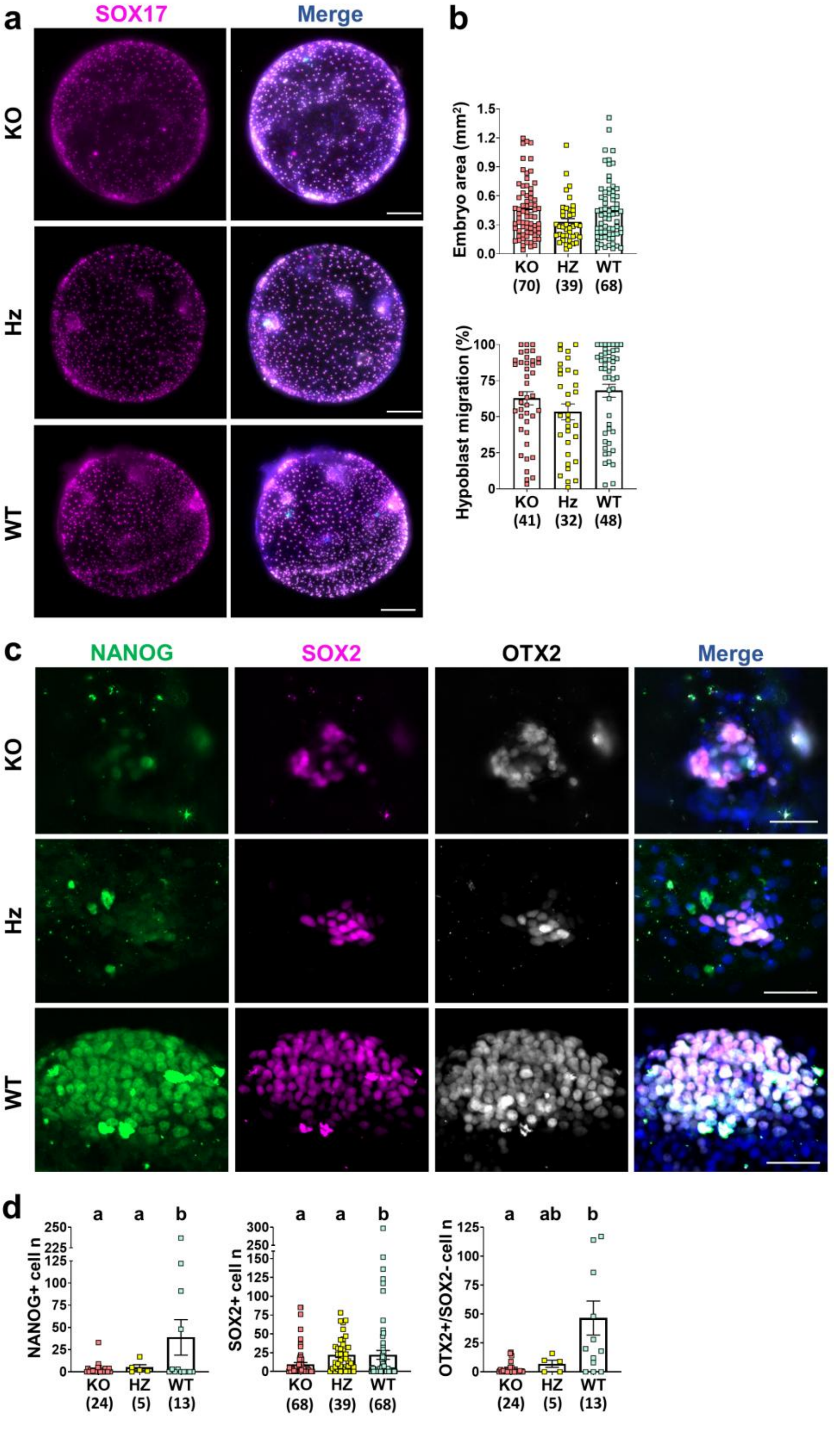


**Supplementary Figure 18.** ***NODAL*** **ablation impairs epiblast survival in day (D) 12** ***in vitro*** **cultured sheep embryos.**

a) Representative immunofluorescence images of *NODAL* KO, Hz and WT D12 *in vitro* embryos stained for SOX17 (magenta; hypoblast). Nuclei were counterstained with DAPI (merge). Scale bars: 200 µm.

b) Scatter plots showing embryo area and the percentage of SOX17+ hypoblast cells migration along the inner embryo surface in KO, Hz, and WT D12 *in vitro* embryos (mean ± s.e.m.). The number of analysed embryos is shown below columns. No statistically significant differences were detected among groups ($P > 0.05$; non-parametric Kruskal-Wallis test).

c) Representative immunofluorescence images of *NODAL* KO, Hz and WT embryonic discs from D12 *in vitro* embryos stained for NANOG (green; epiblast), SOX2 (magenta; epiblast) and OTX2 (white; epiblast and anterior visceral hypoblast). Nuclei were counterstained with DAPI (merge). Scale bars: 50 µm.

d) Scatter plots showing the number of SOX2+ and NANOG+ epiblast cells, and the number of OTX2+/SOX2- anterior visceral hypoblast cells in KO, Hz, and WT D12 *in vitro* embryos (mean ± s.e.m.). The number of analysed embryos is shown below columns. Different letters above columns indicate statistically significant differences ($P \leq 0.05$; non-parametric Kruskal-Wallis test).

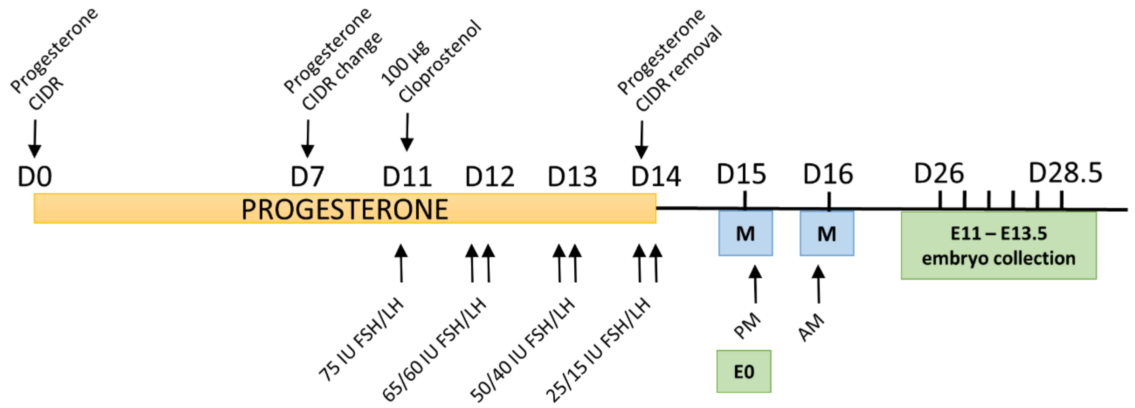


**Supplementary Figure 19. Superovulation protocol employed to obtain *in vivo*-derived sheep embryos**. FSH: Follicle stimulating hormone; LH: luteinizing hormone; IU: international units; M: mating.

**References:**


1 Ramos-Ibeas, P. *et al.* Pluripotency and X chromosome dynamics revealed in pig pre-gastrulating embryos by single cell analysis. *Nat Commun* **10**, 500 (2019). https://doi.org/10.1038/s41467-019-08387-8

2 Simpson, L. *et al.* A single-cell atlas of pig gastrulation as a resource for comparative embryology. *Nat Commun* **15**, 5210 (2024). https://doi.org/10.1038/s41467-024-49407-6

3 Bergmann, S. *et al.* Spatial profiling of early primate gastrulation in utero. *Nature* **609**, 136-143 (2022). https://doi.org/10.1038/s41586-022-04953-1

4 Mayshar, Y. *et al.* Time-aligned hourglass gastrulation models in rabbit and mouse. *Cell* **186**, 2610-2627.e2618 (2023). https://doi.org/10.1016/j.cell.2023.04.037

5 Nowotschin, S. *et al.* The emergent landscape of the mouse gut endoderm at single-cell resolution. *Nature* **569**, 361-367 (2019). https://doi.org/10.1038/s41586-019-1127-1

6 Pijuan-Sala, B. *et al.* A single-cell molecular map of mouse gastrulation and early organogenesis. *Nature* **566**, 490-495 (2019). https://doi.org/10.1038/s41586-019-0933-9

7 Cheng, S. *et al.* Single-Cell RNA-Seq Reveals Cellular Heterogeneity of Pluripotency Transition and X Chromosome Dynamics during Early Mouse Development. *Cell Rep* **26**, 2593-2607.e2593 (2019). https://doi.org/10.1016/j.celrep.2019.02.031

8 Thowfeequ, S. *et al.* An integrated approach identifies the molecular underpinnings of murine anterior visceral endoderm migration. *Dev Cell* **59**, 2347-2363.e2349 (2024). https://doi.org/10.1016/j.devcel.2024.05.014

9 Soneson, C., Love, M. I. & Robinson, M. D. Differential analyses for RNA-seq: transcript-level estimates improve gene-level inferences. *F1000Res* **4**, 1521 (2015). https://doi.org/10.12688/f1000research.7563.2

10 Scatolin, G. N. *et al.* Single-cell transcriptional landscapes of bovine peri-implantation development. *iScience* **27**, 109605 (2024). https://doi.org/10.1016/j.isci.2024.109605

11 Sullivan, D. K. *et al.* kallisto, bustools and kb-python for quantifying bulk, single-cell and single-nucleus RNA-seq. *Nat Protoc* **20**, 587-607 (2025). https://doi.org/10.1038/s41596-024-01057-0
